# Active destabilization of the integron synaptic complex reduces bacterial adaptation to antibiotics

**DOI:** 10.64898/2026.03.10.710562

**Authors:** Ekaterina Vorobevskaia, Pilar Lörzing, Michael Schlierf

## Abstract

Antibiotic multi-resistance (AMR) in bacteria poses a significant threat to global health, driven by mechanisms like the integron genetic system in Gram-negative bacteria. Integrons facilitate AMR by shuffling resistance genes through site-specific recombination, mediated by the integrase enzyme IntI. Earlier studies revealed that the mechanical stability of the synaptic complex, a structure formed by four integrase subunits and DNA, correlates with the recombination efficiency, and, by extension, the adaptation capability. We identified a conserved C-terminal α-helix in IntI that stabilizes the synaptic complex via specific interactions with a binding pocket. To disrupt this interaction, we designed peptides mimicking the α-helix, which reduced the mechanical stability probed with single-molecule optical tweezers. In bacterial adaptation assays, these peptides significantly decreased integron-mediated adaptation to ciprofloxacin stress without exhibiting antimicrobial activity. This approach highlights a novel strategy to combat AMR by targeting integron-mediated gene shuffling, offering potential for future therapeutic development to limit the spread of resistance genes.

## Introduction

Bacteria evolved a multi-functional set of tools for adaptation to challenging environmental conditions, including the presence of antimicrobial agents. Evolution of antibiotic resistance outpaces the developments of new antibiotics since a few decades and poses a great threat to human society (World Health Organization 2014; Murray et al. 2022). In particular, antibiotic multiresistance (AMR) in bacteria leads to rising numbers of AMR-associated deaths and the prognosis is highly alarming (Naghavi et al. 2024). The predominant adaptation tool that promotes AMR in Gram-negative bacteria is the integron genetic system (Davies 1994; Cambray et al. 2010). Integrons are frequently found in different bacterial species ranging from waste water and soil inhabitants to clinical pathogens, and can be both chromosome-associated, e.g. class 4 integrons (Mazel et al. 1998; Rowe-Magnus et al. 2002), or mobile, e.g. class 1 integrons, as a part of a plasmid or a transposon (Sallen et al. 1995; Nandi et al. 2004). Due to their mobile nature, bacterial integrons are taking part in horizontal gene transfer – one of the major mechanisms of AMR transmittance (Stokes and Hall 1989; Davies 1994).

Integrons act as smart and resource-efficient libraries of promotor-less resistance genes that are shuffled upon antibiotic stress in an attempt to position the required gene closest to the cassette promoter (*P*_C_) using site-specific recombination (Fig. 1A) (Cambray et al. 2010). Gene shuffling is mediated by a tyrosine recombinase of the λ phage family called the integrase IntI (Escudero et al. 2015), encoded by an *intI* gene that is being transcribed in the opposite direction of the gene cassettes and under the control of its own promoter (*P*_int_). Both promoters are independent of each other, but are regulated by the transcriptional repressor LexA and activated by the SOS response (Guerin et al. 2009). The bacterial SOS response (Guerin et al. 2009; Spåhr et al. 2018; Michel 2005) itself is activated upon autoproteolytic cleavage of the transcriptional repressor LexA, freeing the operator and making both integron promoters accessible for gene transcription (Collis and Hall 1992). Cassette promoters *P*_C_ were reported of varying strengths with highest expression levels reached for closely positioned cassettes (Jové et al. 2010; Guérin et al. 2011). Between gene cassettes are *attC* sites, which are recognized by the integrase for excision and insertion. They show rather diverse sequences, affecting the DNA recombination efficiency (Loot et al. 2010; Bouvier et al. 2009; Vorobevskaia et al. 2024). The primary gene integration site is called *attI*, located close to the promoter *P*_C_ and shows a rather conserved sequence depending on the class of the integron system (Collis and Hall 2004; Partridge et al. 2000; Hansson et al. 1997; Loot et al. 2024). A series of recombination events, including excision and insertion reactions driven by the recombinase IntI, shuffles genes allowing for a diverse transcriptional landscape within a bacterial population and, thus, leading to a faster, robust adaptation (Escudero et al. 2015).

**Figure 1.**
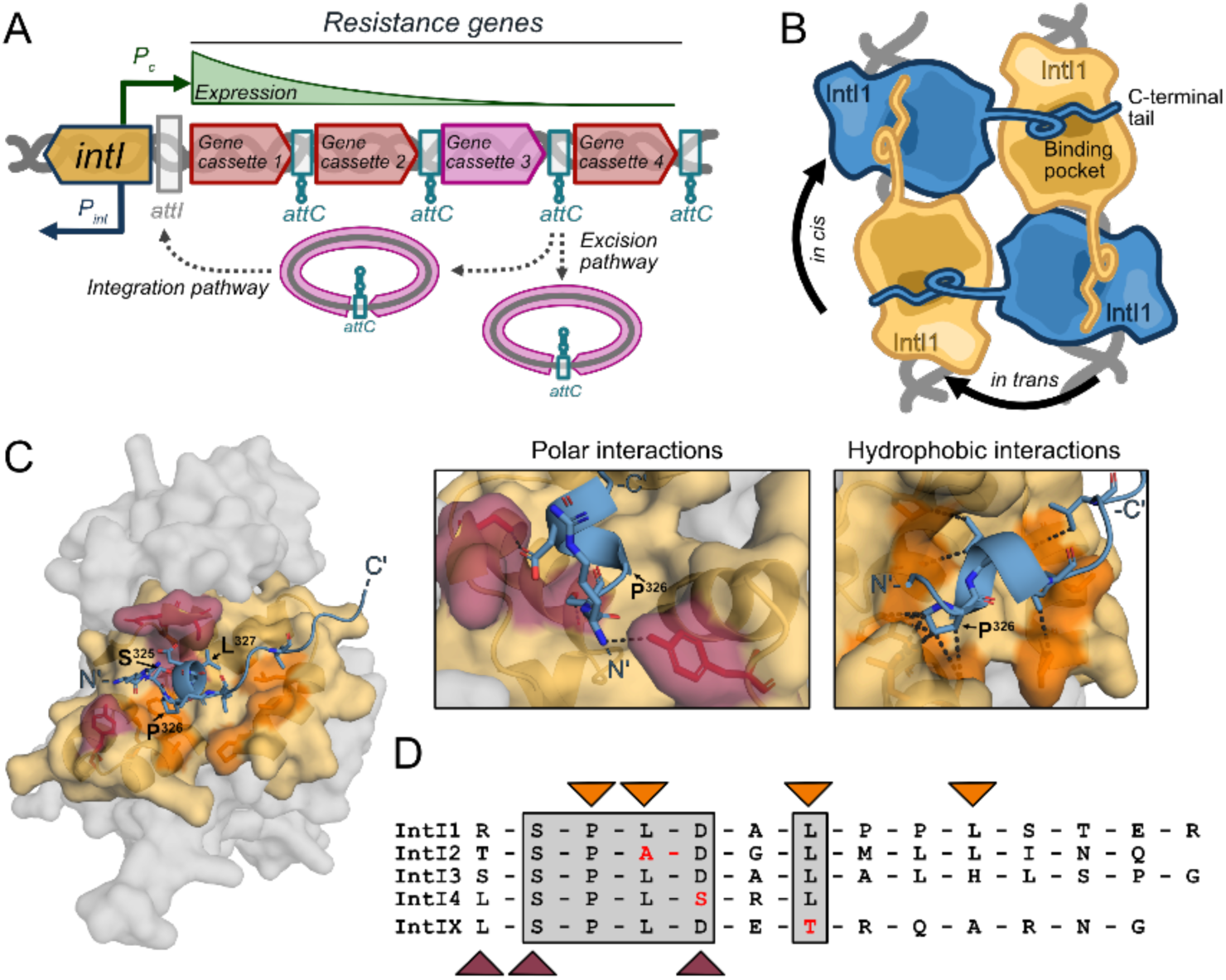
Bacterial integron and integrase structure. **A.** Integron scheme with excision and integration shuffling pathways. **B**. Synaptic complex formation scheme with C-terminal tail loop-like stabilizing docking interaction. Orange IntI represent active integrase subunits, blue IntI, inactive subunits. **C.** Binding pocket and C-terminal α-helix tail interaction of the IntI1 synaptic complex structure predicted by AlphaFold3, zoom-in windows: polar interactions < 3.5 Å; hydrophobic interactions < 4 Å. Visualized with PyMOL (Schrödinger, L., & DeLano, W. 2020). **D.** C-terminal tail sequence alignment across integron integrase classes. Conserved residues are indicated in grey rectangles. Indicated residues: red for polar interactions; orange for hydrophobic contact to the pocket. Sequences were aligned using Clustal Omega Multiple Sequence Alignment (MSA) (Madeira et al. 2024).

Interestingly, the integron system shows a rather high variability in the recombination efficiency that leads to different adaptation rates. Earlier studies have found the *attC* sequence and structure a key determinant for the varying recombination efficiency (Loot et al. 2010; Bouvier et al. 2009; Nivina et al. 2016; Mukhortava et al. 2019). Recently, we could link the recombination efficiency to the mechanical stability of the key assembly of recombination, the so-called synaptic complex (Vorobevskaia et al. 2024). This essential structure is formed by two DNA sites (*attCxattC* or *attCxattI*) and four IntI subunits. Depending on the *attC* sequence and structure, the mechanical stability of the synaptic complex varied and a low recombination efficiency correlated with a low mechanical stability. Less stable complexes are easier to disrupt by forces acting on DNA during processes like transcription, DNA repair, and replication, while more stable synaptic complexes are more likely to initiate recombination.

A crystallographic structure of a class 4 integron synaptic complex also revealed key protein-protein interactions of the macromolecular assembly (MacDonald et al. 2006). Mutating some of these key interactions also lead to a reduced mechanical stability and reduced recombination efficiency (Vorobevskaia et al. 2024). In particular, the C-terminal tail of the integrase caught our attention, as it connects to a binding pocket in the neighboring subunit (Fig. 1B). The C-terminal domain is generally very similar in structure between different tyrosine recombinases and it likely plays a regulating role in recombination (Meinke et al. 2016; Landy 2015; Jayaram et al. 2015). Deletion of the tail in our earlier study, greatly destabilized the complex and diminished bacterial recombination. Thus, this interaction looked particularly promising and could reveal a weak spot to prevent bacterial adaptation or acquisition of new genetic elements using the integron system. While protein engineering allows to understand fundamental mechanisms in general, it does not offer a viable perspective for treatments of AMR patients. Here, we devised a peptide-based strategy to mechanically destabilize the synaptic complex. We designed a set of peptides that could weaken the synaptic complex, dissect key residues of the peptides and characterized their effects on the complex stability. We further wondered if these peptides are also affecting bacterial adaptation and thus developed an adaptation assay allowing us to show that the designed peptide significantly reduces integron mediated adaptation during antibiotic stress, while not showing antimicrobial activity by itself. We anticipate that this idea together with further compound developement could provide a new opportunity to limit effects of AMR and spreading of multi-resistance genes via the bacterial integron system.

## Results

The strong effect of the C-terminal truncation on the mechanical stability and recombination efficiency, in encouraged us to further inspect the tail-mediated inter-integrase interaction. When taking a closer look at the docking interaction between the α-helix and the binding pocket two modes of helix-pocket interactions can be differentiated: *in cis* (active subunit C-terminal α-helix docks to the inactive subunit cavity) and *in trans* (inactive subunit is docking to the active one) (Fig. 1B). The *in cis* interaction mode might add stabilization for the dimerization of integrases on *attC* sites. The *in trans* mode connects both integrase dimers and could be key to stabilize the synaptic complex. To take a closer look at the molecular arrangements, we modeled the IntI1 synaptic complex using Alphafold3 (Abramson et al. 2024) and obtained a structure which is very similar to the IntI4 crystallographic structure (MacDonald et al. 2006). The alignment between both structures showed an RMSD of 2.5 Å (Fig. S1) and like in the IntI4 structure, the Alphafold3 model shows also a binding of the C-terminal α-helix (Cterm) to a binding pocket (BP) on the neighboring subunit (Figure 1C).

Upon closer inspection, we found that the C-terminal tail of IntI1 folds into a small α-helix of 7 amino acids inside the binding pocket, followed by another 7 amino acid-long unstructured chain, which is not present in the IntI4 crystallographic structure. The α-helix forms approximately two turns and is docked in the binding pocket of the receiving IntI1 subunit via a combination of polar and hydrophobic interactions (Fig. 1C and insets). 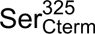 forms likely polar interactions with the backbone of 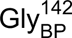, followed by 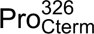 showing several hydrophobic contacts to the pocket with 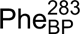, 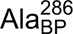 and 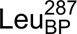 diving into a well-defined dip (Fig. 1C, Fig. S2). The following 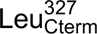 stabilizes the hydrophobic contacts followed by a strong polar interaction between 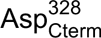 and 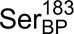. A subsequent bioinformatic analysis revealed that the SPLD motif at the C-terminal end is highly conserved among integron integrases (Fig. 1D). Interestingly, most C-terminal tails of integrases are longer, than the structured α-helix and have an additional, likely unstructured chain, like IntI1.

Building on the stabilizing role of the C-terminal tail docking and the high conservation of the C-terminal tail among integrases, we hypothesized that blocking the binding pocket would destabilize the synaptic complex and, by extension, reduce bacterial adaptation (Fig. 2A). We introduce a new approach of ***active synaptic complex destabilization*** of integrases by applying a small, α-helix-like molecule, blocking the integrase binding pocket, while being independent of the *attC* sites and the integrase class.

**Figure 2.**
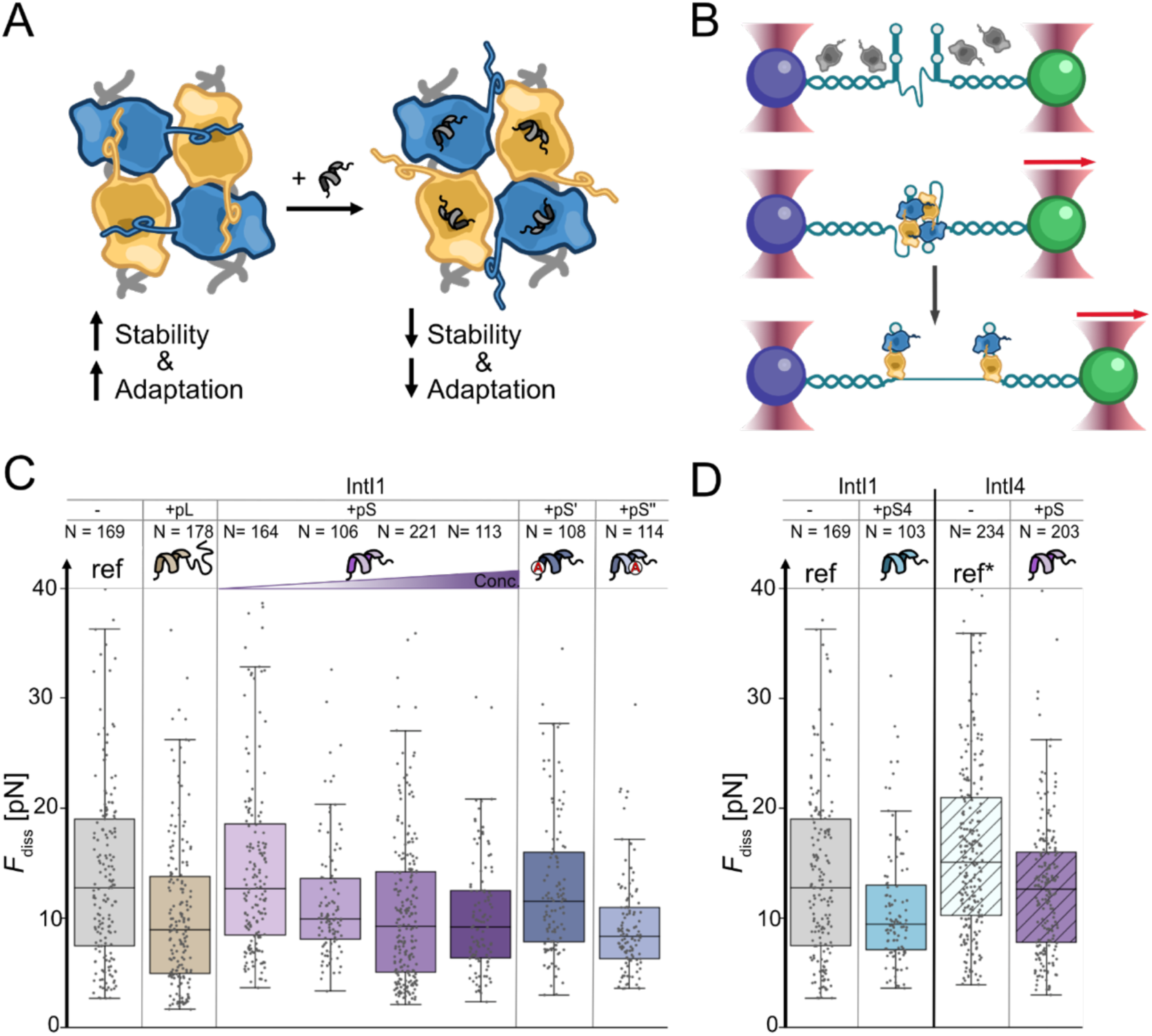
Active synaptic complex destabilization and mechanical stability. **A.** Active synaptic complex destabilization concept: addition of a small peptide prevents C-terminal tail loop-like interaction. **B.** Force spectroscopy stability assay using optical tweezers; synaptic complex disassembly force indicates mechanical stability. **C.** Characteristic synaptic complex disassembly forces *F*diss as a measure of mechanical stability for the reference synaptic complex IntI1-*attCaadA7*^bs^ (“ref”) and in the presence of different peptides (N - number of measured disassembly events; concentration of pL, pS’ and pS’’ was 10 µM). pS peptide was measured at increasing concentrations (2 µM, 5 µM, 10 µM and 50 µM) as indicated by the color gradient ramp. **D.** Cross-class destabilization activity of mimicking peptides from IntI4 C-terminal α-helix (pS4) on IntI1-*attCaadA7*^bs^ (“ref”) synaptic complex and of mimicking peptides from IntI1 C-terminal α-helix (pS) on IntI4-*attCaadA7*^bs^ (“ref*”) synaptic complex. Concentration of pS4 and pS was 10 µM.

As a starting design for the small competing molecule, we designed a peptide based on the highly conserved regions of the C-terminal of integron integrases. The first design was a 14 aa-long peptide (pL), with the sequence of the IntI1 C-terminus encompassing the short α-helix followed by the unstructured region. To test the influence of the peptide on the synaptic complex of four integrase subunits and two *attC* sites, we have adapted our earlier single-molecule force spectroscopy assay in optical tweezers (Fig. 2B) (Vorobevskaia et al. 2024). In short, using an optical tweezers setup we tethered a DNA substrate containing long double-stranded DNA handles and a mimicry of an integron cassette, consisting of two *attC* sites and a single-stranded DNA spacer between two microspheres (Fig. S3). At low forces, both attC sites can fold into their characteristic secondary structure (Mukhortava et al. 2019) and in presence of integrase a synaptic complex can be self-assembled. Afterwards, we increase the force on the complex, which eventually disassembles showing a characteristic fingerprint which allows us to identify that a synaptic complex was present before (Fig. S4). The characteristic disassembly force of the synaptic complex is the measure of its mechanical stability. As a reference, we use the well-studied, highly stable IntI1-*attC_aadA7_*^bs^ synaptic complex in absence of the peptide (Fig. 2C, Table S1). Through our microfluidic system we introduced pL and the synaptic complex self-assembled in its presence. Subsequently, we found a significant decrease of the median disassembly force from *F*_dis_ = 12.7 pN to *F*_dis_(pL)= 8.9 pN suggesting a specific and strong interaction between pL and the integrase monomers, outcompeting the natural C-terminal tails of the integrases themselves.

Upon a closer inspection of the homology sequence and the structure of the IntI4 synaptic complex (MacDonald et al. 2006), we wondered if a minimal peptide motif mimicking the bound α-helix would be sufficient to destabilize the synaptic complex. Therefore, we decided for a 7 amino acid short peptide (pS) based on the Alphafold-predicted α-helix structure of IntI1. Upon titration of pS, we observed a distinct dose dependent reduction of the median disassembly force (Fig. 2C). At [pS] = 2 µM, we did not observe a significant decrease of *F*_dis_, while at 5 µM *F*_dis_ dropped significantly and plateaued at 10 µM at *F*_dis_ ∼ 9.2 pN (Fig. 2C). Concentrations up to 50 µM did not show any further destabilization. We conclude, that very likely the short peptide is sufficient to specifically occupy the binding pockets and prevent a mechanically stable synaptic complex.

### Single amino acid substitutions in pS modulate its efficacy

pS and pL are mimicking the IntI1 C-terminal tail sequence, however even the conserved core residues of the α-helix have sequence variations in other integron integrase classes. Therefore, we designed two variants of pS introducing Alanine substitutions based on our structural analysis of the Alphafold3 predicted IntI1 synaptic complex. In pS’ Ala substitutes Ser^2^ potentially breaking a strong polar interaction to the binding pocket (Fig. 1C and 1D). Notably, Ser^2^ is conserved in all known integrases, suggesting that a substitution might have severe impact. In pS’’ we substituted Leu^4^ with Ala, where the Leu showed a deep dive into the binding pocket, but at the same time it is not as conserved (Fig. 1C and 1D). In the optical tweezers mechanical stability assay, we found that [pS’] = 10 µM lost the destabilization ability and *F*_dis_ (pS’) = 11.5 pN remained nearly unchanged compared to an experiment in absence of pS’ the undisturbed complex (Fig. 2C). The loss of destabilization for pS’ indicates that the Ser^2^-mediated polar interaction to the pocket is vital for the stable docking of the peptide, in good agreement with the strict conservation among integron integrases. pS’’, instead, showed at 10 µM synaptic complex destabilization. The median disassembly force even dropped slightly more (8.3 pN, N=114) compared to pS, suggesting a slightly increased efficacy. Notably, the Leu^4^ to Ala substitution does not change the chemical nature of the hydrophobic side chain, though is less bulky, and thus might facilitate a tighter contact to the surface of the binding pocket or an increased binding rate.

### Cross-class destabilization activity of the peptides on integrases

Both pL and pS peptides were able to destabilize the synaptic complex of IntI1. However, the variants indicated that some sequence flexibility might be tolerated, thus we wondered if it would be possible to dock a peptide mimicking the α-helix of another integron class to the pocket of IntI1? We tested this by designing a short peptide based on the C-terminal tail of IntI4 (pS4) (Fig. 1D). We preformed our stability assay with the reference IntI1-*attC_aadA7_*^bs^ synaptic complex and saw the same level of destabilization as for pS (Fig. 2D). The median disassembly force of the IntI1-*attC_aadA7_*^bs^:pS4 complex was *F*_dis_ (pS4^10^ ^µM^) ∼ 9.4 pN (N = 103) and the rather narrow distribution of disassembly forces, similar to pS^50^ ^µM^ suggests potentially an increased binding despite the large aspartic acid side chain instead of a serine side chain. Encouraged by the activity of pS4 on the destabilization of the IntI1 synaptic complex, we wondered whether pS exhibits also cross-class activity on synaptic complexes by another integrases. Therefore, we recombinantly produced IntI4, an integrase class found on chromosomal integrons, e.g. in *Vibrio cholerae*. We found that IntI4 forms like IntI1 mechanically stable synaptic complexes even with the atypical *attC_aadA7_*^bs^. Interestingly, the IntI4 synaptic complex showed even a slightly increased stability compared to IntI1 (Fig. 2D). This complex was readily destabilized by addition of pS from *F*_dis_ (IntI4) ∼ 15.0 pN (N = 234) to *F*_dis_ (IntI4 + pS^10^ ^µM^) ∼ 12.6 pN (N = 203), illustrating that the designed peptide could potentially also be active in evolutionary related integron systems.

### Integron-mediated bacterial adaptation to ciprofloxacin is reduced by pS

Thus, we wondered if such a destabilization also leads to reduced integron-mediated adaptation in bacteria. To this end we mimicked an adaptation process with a designed low copy plasmid carrying a bacterial integron system transformed into *E. coli* MG1655 (strain #965). The plasmid carries a permanent kanamycin resistance gene for selection and a class 1 integron with *intI1* and a library of four antibiotic resistance gene cassettes containing the open reading frames for *smR2* (*e.g.* Streptomycin resistance), *catB3* (*e.g.* Chloramphenicol resistance), *blaVIM4* (*e.g.* Penicillin resistance), and *aac(6’)-lb-cr* (*e.g.* Ciprofloxacin resistance) (Fig. 3A, Figure S5). IntI1 is expressed from its native SOS-regulated promoter without overexpression, the cassettes are expressed under the control of *P*_cw_, thus, mimicking a naturally occurring mobile integron plasmid. To induce antibiotic stress, we have selected ciprofloxacin because it also directly activates the SOS response in *E. coli* MG1655 (Phillips et al. 1987; Vikedal et al. 2025; Teichmann et al. 2025) and, by extension, the integron system (Mazel et al. 1998). The corresponding ciprofloxacin resistance gene is located at the end of our designed gene cassette library of the integron, which makes its expression extremely weak and strengthens the pressure on integron adaptation. To assess the efficiency of our peptide against adaptation, we have devised several assays and monitored multiple survival parameters of the bacterial culture. At first, we devised a strategy how to provide the peptide to the cytosol of bacteria via chromosomal expression (Fig. 3B). To this extent, we inserted into MG1655 an *mCherry-pS* fusion in the *lac* operon using λ Red recombineering (Fig. S6), such that we could induce expression by addition of IPTG (Schärfen et al. 2020). We transformed further the integron plasmid to this strain and obtained *E. coli* MG1655 strain #1008. Using fluorescence microscopy, we confirmed that in absence of IPTG, the mCherry-pS peptide was not significantly expressed. We monitored cell growth of strains #965 and #1008 and did not observe a change in growth rate, average cell length, nor a change in cell density after 19 hours of growth in LB_kana_ (Fig. 3B, Table S2). Though, upon IPTG induction (#1008 in presence of IPTG; thereafter #1008_IPTG_), we observed a strong expression of mCherry-pS (Fig. 3B #1008_IPTG_ and Fig. S7), which was absent in #965 and barely detected in #1008 in absence of IPTG, indicating minimal basal expression from *P*_lac_. Even under expression of mCherry-pS, bacterial growth did not show any significant changes or deleterious effects on bacterial growth, thus mCherry-pS does not exhibit any bactericidal activity itself.

**Figure 3.**
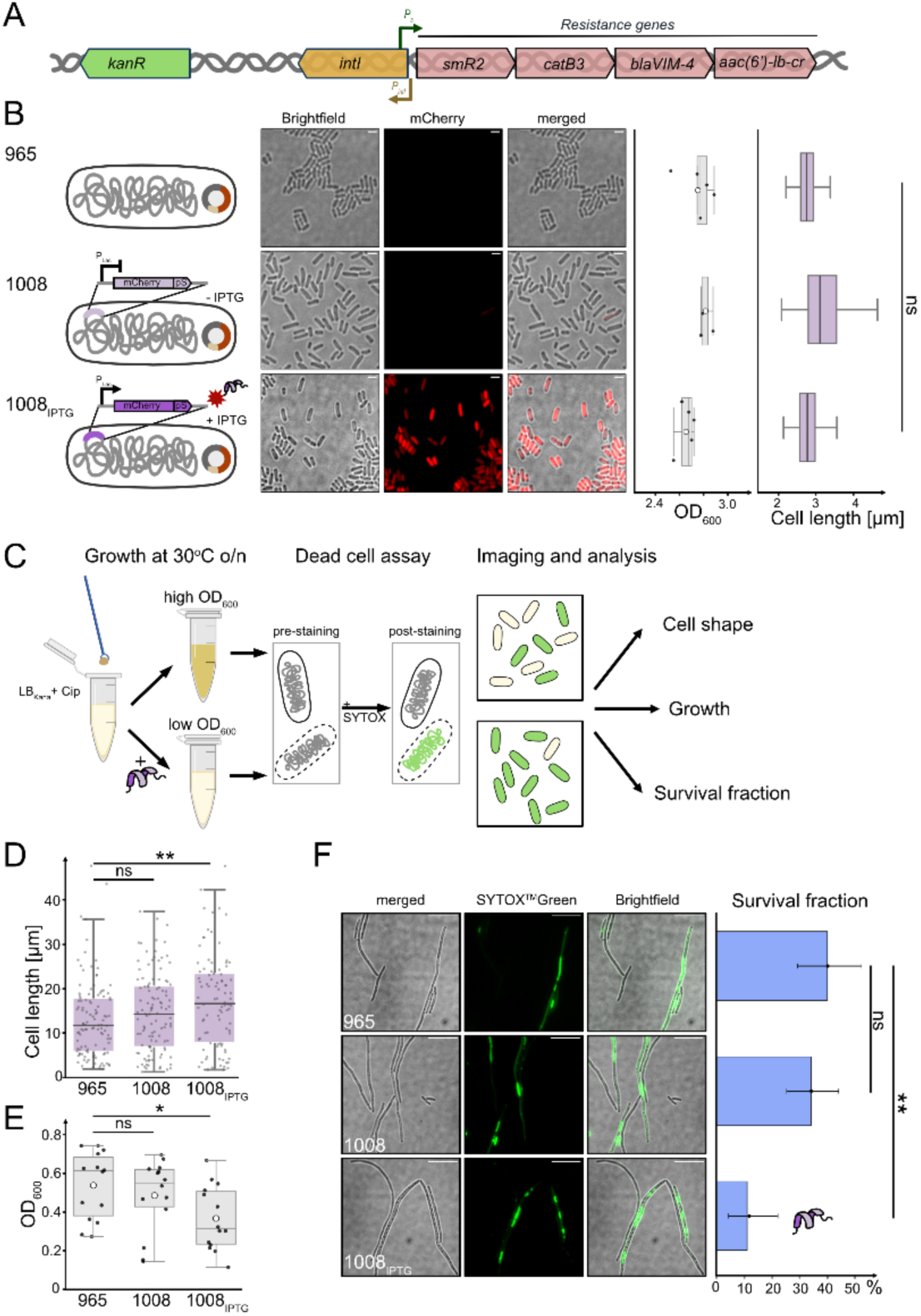
Peptide reduces adaptation and survival in bacteria. **A.** Schematic representation of the p15A-integron plasmid. **B.** pS is introduced in the chromosome as a C-terminal fusion to mCherry in the *lac* operon under the control of the *lac* promoter (strain #1008). Expression of mCherry-pS is induced with 1 mM IPTG (#1008IPTG) and is absent without IPTG (#1008). The strain lacking the chromosomal mCherry-pS construct (#965) shows no detectable background fluorescence. mCherry images are displayed using identical intensity scaling to allow direct comparison between strains. OD600 values after 19 h growth and single cell length for #965, #1008 and #1008IPTG do not vary significantly (Welch t-test, OD600 p-value = 0.19; cell length p-value =0.2). Scale bar 2 µm. **C.** Scheme of the adaptation assay to determine the effects of pS on bacterial adaptation to ciprofloxacin stress in liquid culture. LBKana-Cip medium is inoculated with the bacterial strain and grown overnight at 30°C with continuous shaking and subsequent OD600 measurement. Resulting cultures were stained with SYTOX Green for the dead cell assay and the survival fraction determined using single-cell image analysis. **D.** Cell length of #965, #1008, and #1008IPTG after 19 h Ciprofloxacin stress leads to cell elongation. Cell length in presence of mCherry-pS is significantly longer than in absence. **E.** OD600 after 19 h Ciprofloxacin stress of #965, #1008, and #1008IPTG; white circles depict the mean. **F.** Dead cell assay imaging results for studied strains. SYTOX Green signal in the cell indicates cell death. Images are displayed using individually adjusted contrast settings for visualization purposes, quantitative analysis was performed on the original, unprocessed data. Scale bar 10 µm. Bar plot shows the survival fraction of the imaged #965, #1008, and #1008IPTG. Survival fraction in presence of mCherry-pS is significantly shorter.

Subsequently, we devised an adaptation assay that induces antibiotic stress on our engineered *E. coli* strains (Fig. 3C). For this, bacteria were inoculated and grown over night in LB_Kana_ in presence of 40 ng/µL ciprofloxacin (1.3x MIC) to trigger the integron-mediated adaptation (Figure S8). During the overnight growth we added IPTG to distinct cultures, to induce mCherry-pS production, thus having pS present to potentially inhibit integron-mediated adaptation. After the overnight growth, we have measured the apparent cell density (OD_600_), used brightfield microscopy to determine cell shape and length and performed a fluorescence based dead cell assay to monitor the survival fraction of cells.

As expected for fluoroquinolone treatment, all strains triggered the SOS response, which lead to a filamentous morphology as compared to the typical rod-shaped morphology of unstressed *E. coli* due to SulA expression (Aldred et al. 2014; Phillips et al. 1987). Cell elongation was heterogeneous between cells and between strains (Fig. 3D). Without mCherry-pS expression the length increased by ∼ 5-6 times compared to the length of the unstressed cells reaching a mean cell length of 13-14 µm. Cells grown in presence of IPTG (#1008_IPTG_) showed an even stronger heterogeneity in cell length and a mean cell length of 17 µm. As a control, we provided generated also a strain carrying an integron plasmid with an IntI1^ΔC^ variant (#1016), which showed even lower mechanical stability and low recombination efficiency in earlier studies (Vorobevskaia et al. 2024). This strain exhibited an even slightly higher cell length and increased heterogeneity phenotype (Fig. S9, Table S2). As an independent measure of cell growth, we determined also the apparent cell density by determining the OD_600_ after 19h growth. While cell length indicated already an effect of the mCherry-pS overexpression, the OD_600_ showed an even stronger effect. Both #965 and #1008 showed after 19h growth an OD_600_∼ 0.6, suggesting despite the strong antibiotic exposure with [CIP] = 1.3x MIC, some bacteria were able to adapt and continued growing in contrast to MG1655 in absence of the integron system (Fig. S8). Upon IPTG induction, and, by extension, the presence of mCherry-pS, the mean OD_600_ dropped to 0.3, showing a significantly decreased cell growth (p value = 0.013 according to Welch t-test) (Fig. 3E). Taken together the increase of the mean cell length and a reduced optical density, this change is indicating that ciprofloxacin impacted the growth parameters to a greater degree than for the strains without peptide expression, suggesting that mCherry-pS suppresses bacterial adaptation. Yet, both cell length and optical density are only indirect measures of cell survival.

To assess the survival population after the overnight ciprofloxacin stress, we have implemented a fluorescent dead cell assay where cells are stained with SYTOX green prior to fluorescence imaging (Fig. 3C). SYTOX green is a membrane impermeable fluorophore, which stains DNA. Since ciprofloxacin does not directly impair the bacterial membrane, viable cells will not get stained with SYTOX, while dead bacterial cells likely lost their fully intact membrane and we will observe direct staining of the remaining chromosome. We have subsequently determined the fraction of unstained cells as surviving cells and termed this the survival fraction. In absence of ciprofloxacin, we found a 98% survival fraction after overnight growth, while upon ciprofloxacin stress both strains #965 and #1008 showed only ∼ 40% survival fraction. Expression of mCherry-pS (#1008_IPTG_) reduced the survival fraction 4-fold to ∼ 11%, hence in presence of pS. Hence the integron enabled every second-to-third bacterial cell to survive after ciprofloxacin treatment, the addition of pS reduced this chance to only one out of ten cells. Noteworthy, SYTOX staining combined with single-cell microscopy further revealed a very heterogenous staining pattern: some cells showed homogeneous fluorescence throughout the cell, while others showed fluorescence patches, suggesting separated condensed chromosomal DNA, a phenotype recently reported in response to ciprofloxacin (Bos et al. 2015; Vikedal et al. 2025) (Fig. 3F).

### pS reduces shuffling of the integron system

pS is originally designed to destabilize the synaptic complex, the key complex during DNA recombination, the essential step in integron-mediated adaptation. Thus, since mCherry-pS significantly reduces the survival fraction of cells, it should also decrease the degree of cassette shuffling. Cassette shuffling or cassette excision is one mechanism of the integron to activate “dormant” resistance genes, far away from the cassette promoter *P*_cw_. Therefore, we aimed to detect the degree of cassette shuffling occurring after overnight ciprofloxacin stress. Taking into account the initial composition of the integron cassette library on the plasmid, we have designed a pair of PCR primers that produce the amplicon between the cassette promoter until the end of the *aac(6’)-ib-cr* gene cassette, conferring resistance to ciprofloxacin (Fig. 4A). Without shuffling, the *aac(6’)-ib-cr* gene cassette is located on the 4^th^ position, furthest away from the promoter and the amplicon size is ∼ 3200 bp. However, if shuffling occurred and the gene cassette was relocated closer to the promoter, the size of the amplicon would be reduced. This means, if the *aac(6’)-ib-cr* gene appears at the third cassette position, amplicon size is estimated to be ∼ 2500 bp, for the second position ∼ 1600 bp, and for the first position ∼ 700 bp. Instead of colony PCR reaction, we devised a protocol to use directly the liquid overnight culture as PCR template (see Materials and Methods) in order to maximize the range of sampled cells. The resulting PCR product was subjected to electrophoresis on an agarose gel (Fig. 4B). The most dominant band for all strains was the original 4^th^ cassette position, indicated by an intense band on the gel at the level of ∼3200 bp (Fig. 4B). However, upon Ciprofloxacin stress, we observed also shorter amplicons in strain #965 at the expected sizes, which likely originate from an active integron system shuffling the gene cassettes resulting in *aac(6’)-ib-cr* to be relocated to all possible cassette positions. To demonstrate that the smaller amplicons result from specific integron-shuffling, we have tested a strain with a catalytically inactive IntI1_Y312F_ and observed only an amplicon at the original 4^th^ cassette position. Additionally, in absence of ciprofloxacin stress, we did also observe only the 4^th^ cassette position amplicon, in good agreement with earlier findings that the integron system is activated by stress leading to the SOS response (Guerin et al. 2009). Next, we wondered, if mCherry-pS suppresses gene shuffling of the integron system. Therefore, we tested the strain #1008_IPTG_, which produced mCherry-pS, we did observe mainly amplicons at the 4^th^ cassette position, with very faint bands corresponding to 3^rd^ position. This indicates that shuffling, though still being present, was strongly reduced or less successful. Thus, together with the reduced growth phenotype, the significantly reduced survival fraction and reduced shuffling efficiency, we suggest that pS successfully inhibits the integron function and weakens its adaptation ability.

**Figure 4.**
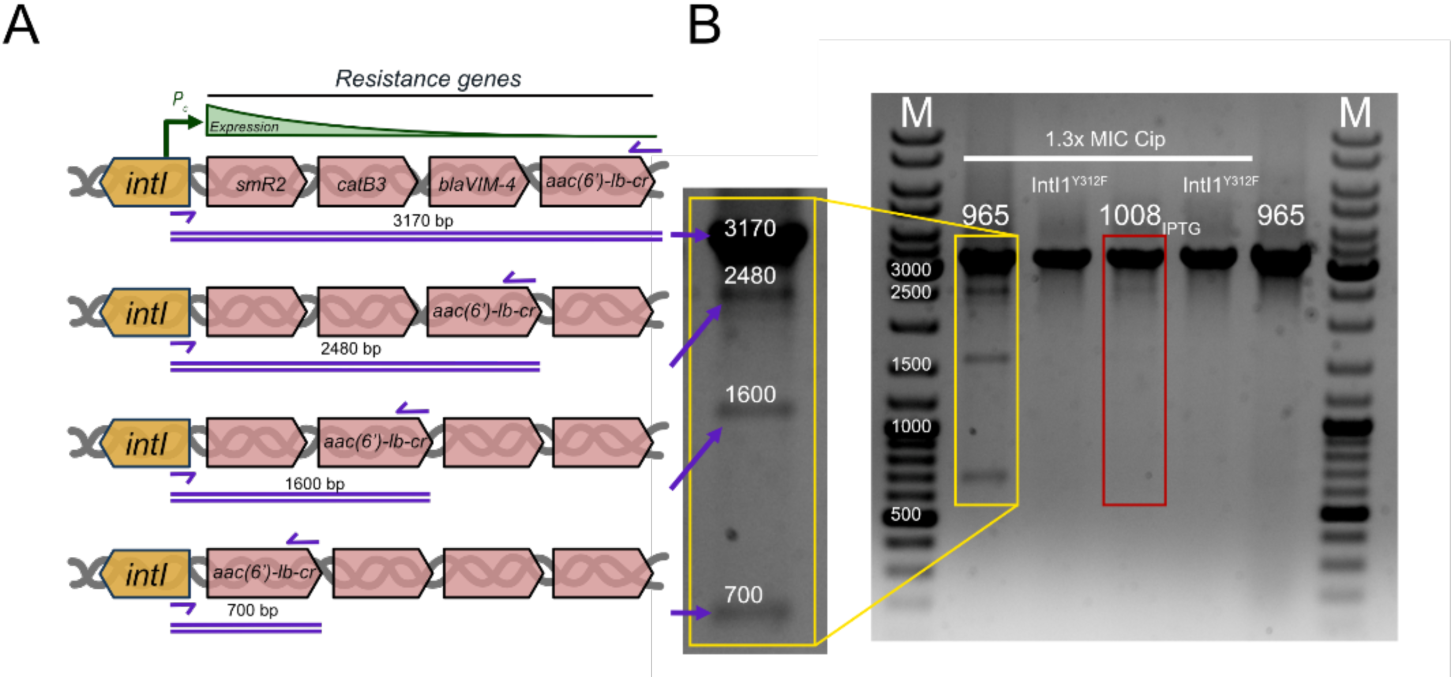
Ciprofloxacin stress adaptation. **A.** Schematic representation of the ciprofloxacin resistance gene cassette (*aac(6’)-lb-cr*) shuffling position possibilities in the integron gene cassette library of the studied strains. Cassette position was determined through PCR using a pair of primers indicated as violet arrows, expected PCR product sizes are denoted. **B**. Agarose gel of PCR product to identify cassette positions. DNA ladder is indicated with “M”. The studied strains are indicated, including controls: IntI1Y312F corresponds to the catalytically inactive integrase protein that is unable to perform recombination and shuffle cassettes. The last lane is culture control without Ciprofloxacin stress.

## Discussion

Multi-resistant bacteria are a major concern, also due to the lack of discovery of novel antibiotics and the rapid evolution of resistances (Murray et al. 2022). The bacterial integron is the predominant system for adaptation in Gram-negative bacteria and carries frequently multiple antibiotic resistance genes (Mazel 2006; Moura et al. 2009). In earlier studies, an Achilles heel of a key step in DNA recombination of the integron system, namely the interaction between the C-terminal extension of the recombinase IntI1 and a binding pocket on the neighboring recombinase unit, was identified (Vorobevskaia et al. 2024). A deletion of this extension destabilized the synaptic complex and reduced the recombination efficiency significantly. Yet, recombinase mutations are impractical for potential future patient treatment. Here, we developed an active synaptic complex destabilization approach to decrease bacterial adaptation to antibiotic stress by blocking the C-terminal tail interaction using designed peptides. We have analyzed the sequence and structure of C-terminal tails of different classes of integrases and identified a common and highly conserved **SP^326^L**(D) motif that is predicted to form a short α-helix interacting with the binding pocket through both polar and hydrophobic interactions (Fig. 1D). For instance, Pro^326^ has a capacity to form a multitude of hydrophobic interactions as well as has a geometrically precise fit into a distinct cavity in the binding pocket. More generally, a C-terminal tail α-helix is present also in other tyrosine recombinases. For instance, *E. coli* recombinases XerD, XerC, and an integrase from a bacteriophage HP1 share a short terminal **SPLS** sequence (Ashkenazy et al. 2016; Subramanya et al. 1997; Castillo et al. 2017; Hickman et al. 1997) and also form synaptic complexes during recombination. Beyond that also for biotechnologically used recombinases like Cre or Flp, a C-terminal connecting α-helix was reported (Gopaul et al. 1998; Y. Chen et al. 2000). This general pattern of the C-terminal α-helix function might imply an evolutionary-determined interaction mode that adds synaptic complex stability and promotes recombination efficiency. The α-helical structure of the core part of the C-terminal tail is likely necessary for tight interactions into the designated binding pocket, however, most integrases have also an unstructured extension beyond the short α-helix. For IntI1, this extension is seven amino acids long and has according to the Alphafold-predicted structure one potential hydrophobic interaction to the edge of the binding pocket. Therefore, as an initial candidate for the active synaptic complex destabilization, we have designed the 14 aa long pL peptide that exactly mimics the C-terminal tail of the IntI1 integrase, including the unstructured chain, which we omitted in the second 7 aa long pS design. Both, the 14 aa and the 7 aa long peptides, mechanically destabilized in a dose-dependent fashion the synaptic complex. The overall distribution of disassembly forces, yet, remained quite broad, with occasional disassembly events occurring above 25 pN resembling the behavior of the reference IntI1-*attC_aadA7_*^bs^ complex without peptide interference. This might indicate insufficient saturation of the IntI1 monomers with the peptides, higher unbinding rates and a strong out-competition by the existing C-terminal tail in IntI1. This calls for further optimization of the compound, lower unbinding rates, higher binding rates and thus increased overall efficacy. From a mechanistic point of view, through single point mutations we could abolish (pS’) or strengthen (pS’’) the destabilization activity on the synaptic complex. Such site-specific interaction modifications open possibilities to further peptidomimetic optimizations – by using structural data of the peptide-pocket interaction and strengthening the important contacts between residues.

Interestingly, the initial design was based on class 1 integrons, though we discovered cross-class activity, likely due to the high sequence conservation between integron classes. In particular, Asp^5^Ser substitution is not impairing the activity of the peptide and pS4 could destabilize the IntI1 synaptic complex. The question of cross-class activity is not only of basic research interest, but also in the view of potential health care applications of the active synaptic complex disassembly approach. It is important that a destabilizing molecule shows activity on different classes of integrases as it is challenging to determine integron classes in a timely manner. While class 1 integrons are clinically the most relevant adaptation systems (Rowe-Magnus et al. 2002; Davies 1994; D.-Q. Chen et al. 2018; Escudero et al. 2015), chromosomal class 4 integrons from e.g. *Vibrio cholera*, were recently discovered to carry anti-phage genes (Darracq et al. 2025; Kieffer et al. 2025; Richard et al. 2022), which could become relevant in the rising field of phage therapy.

Yet, it remained unclear if the mechanical destabilization can be directly related to a reduced bacterial adaptation capacity. Here, we did not observe any effects on bacterial growth during overexpression of the peptide, suggesting that the peptide itself does not exhibit antimicrobial activity. Thus, it might not act as selection pressure for an evolutionary escape (Darby et al. 2023; Santajit and Indrawattana 2016). Yet, we could directly show that the peptide successfully reduced integron-mediated bacterial adaptation to antibiotic stress due to lower survival fraction and reduced gene shuffling. In particular, the reduction in the survival fraction is of interest – as these are the cells that have survived the antibiotic stress and can potentially restore the population after the stress is removed – the core of the resistance population. If we think about a translational context then each antibiotic-surviving bacterium is a potential threat to the patient’s health and the lower the fraction of these cells is the better for the outcome of the patient treatment. Noteworthy, the peptide showed high activity in bacteria despite the relatively large fluorescent protein fusion. This suggests that further improvement and new strategies for peptide delivery to Gram-negative bacteria is of high interest for further development. Overall, we anticipate that further peptidomimetic optimization of pS could open up new avenues to reduce the integron-mediated spread of antibiotic resistances and adaptation to antibiotic stress.

Furthermore, we anticipate that this integron-specific discovery could be expanded to multi-protein complexes from either the tyrosine-recombinase family or other systems involving protein-protein contacts mediated by a small peptide-binding pocket interaction between partners.

## Materials and Methods

### Optical tweezers double-attC DNA construct

To assemble the reference IntI1-*attC_aadA7_*^bs^ synaptic complex *in vitro* for the optical tweezers experiments, we have used an engineered DNA construct described previously (Vorobevskaia et al. 2024). In brief, it consists of a ssDNA insert with two *attC_aadA7_*^bs^ hairpins and a spacer, which provides a molecular fingerprint to identify synaptic complex disassembly, and of two dsDNA handles (each 2.5 kbp length) for tethering to the beads for optical tweezers experiments (Fig. S3). Instead of assembling the molecule from three separate parts via ligation, we have constructed a new plasmid that carries the both DNA handle sequences flanking the DNA region to be converted to ssDNA using In-Fusion assembly (Takara Bio, Japan). The DNA construct with both DNA handles and the insert (5256 bp in length) was produced by PCR using primers carrying either a 5’ biotin or 5’ triple-digoxigenin modification for specific attachment to the beads (primers listed in Table S3). Based on earlier developments to produce distinct ssDNA regions (Belan et al. 2021; Taylor et al. 2022) we introduced two specific nicking sites for the Nb. BbvCI (New England Biolabs, USA) to provide nicks on the double-stranded DNA. The PCR product was nicked, cleaned using the DNA cleaning kit (Machery-Nagel, Germany) and stored for optical tweezers experiments at 4°C. Full sequence details are provided in (Table S3).

### Single-molecule optical tweezers experiments with the peptides

Force spectroscopy measurements were performed on a high-resolution optical tweezers instrument (C-trap, LUMICKS, Netherlands). The traps were calibrated to have a stiffness of 0.4-0.5 pN/nm. First, the nicked DNA construct was incubated with 2.11 µm anti-digoxigenin coated polystyrene beads (0.1% w/v, Biozol, Germany) for 15 min at room temperature. The construct was then tethered to the 1.76 µm streptavidin coated polystyrene bead (1% w/v, Spherotech, USA) using laminar flow of the microfluidic chamber (Fig. S3). The established tether was overstretched without flow, by moving one trap away and the melting DNA was confirmed using the real-time force-extension curve analysis. The overstretching at the force of ∼60 pN produced a plateau with uneven unfolding events corresponding to the dsDNA melting process during which 256 bp of the nicked strain was peeled away, leaving the ssDNA insert sequence, similar as described earlier (Belan et al. 2021). The tether was subsequently relaxed and compared to the expected calculated contour length of the tether (∼5.2 kbp dsDNA ≈ 1.76 µm) as well as the steepness of the force-extension slope to verify successful hybrid dsDNA-ssDNA generation. Afterward, the tether was moved to channel 4 of the microfluidic chamber that contained measuring buffer with 120 nM of integrase (IntI1 or IntI4) and 10 µM peptide (except in cases of pS peptide titration experiment – there 2, 5 and 50 µM were used as well) (full protein and peptide sequences in Table S4). The relaxed tether was then allowed to form a synaptic complex and the measurements were conducted as described previously (Vorobevskaia et al. 2024). The measurements were performed in PBS buffer (Gibco, USA) with the addition of enzymatic POC oxygen scavenger system (Swoboda et al. 2012) at a constant pulling velocity of 250 nm/s.

### Force-extension curve fitting and analysis

Force-extension curves were used to characterize the tether’s behavior during stretch-and-relax cycles. These curves were analyzed using a hybrid worm-like chain (WLC) polymer elasticity model, which is based on Marko-Siggia’s WLC model (Mukhortava et al. 2019) and incorporates enthalpic stretching modifications (Wang et al. 1997). When a synaptic complex forms, the applied force primarily acts on the dsDNA and the rigid synaptic complex. Thus, the force-extension behavior was initially modeled using the extensible WLC model (Eq. 1) (Wang et al. 1997; Bustamante and Yan 2022):

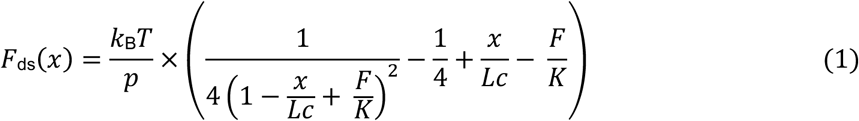

Here, *p* is the persistence length, *K* is the stretch modulus, *L*c is the contour length of dsDNA, and *x* is its extension. *k*_B_ is the Boltzmann constant, and T is set to room temperature.

After synaptic complex disassembly, the construct includes a flexible 114-nt-long ssDNA, requiring a hybrid WLC (hWLC) model to account for the differing flexibility of dsDNA and ssDNA (Mukhortava et al. 2019). The ssDNA term (Eq. 2) uses p_ss = 2 nm, L_ss as the ssDNA contour length, and x_ss as its extension:

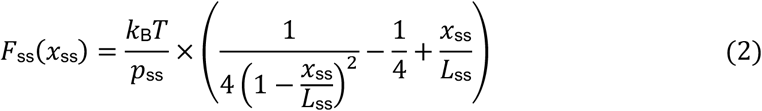

The total construct extension was calculated after WLC inversion using Cardano’s approximation:

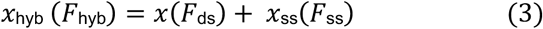

The hybrid force-extension behavior was fitted to the data, and the contour length change from the initial dsDNA handle to the hybrid WLC construct of ∼ 71 nm served as a molecular fingerprint to confirm complete synaptic complex disassembly.

### Disassembly and Unfolding Force Analysis

Data analysis was performed in Python 3.10.5 using Jupyter Notebook and standard Python packages (Table S5). The lumicks.pylake library was used to extract and analyze data from the C-trap. Here, high-frequency force and piezo distance data (78 kHz) were downsampled by a factor of 300. Force-extension curves (FECs) were plotted using the force signal from the immobile Trap 2 against the extension (piezo distance) and corrected for trap crosstalk. Measurements included pulling and relaxing cycles, reaching at least 30 pN force to ensure full disassembly and hairpin unfolding. Relaxation traces, representing the most unfolded state in the presence of the protein, were used to classify pulling traces based on observed integrase events using hWLC fitting and molecular fingerprints. Synaptic complex disassembly events were identified as peaks in the FECs, indicating transitions from the synaptic complex to integrase-bound hairpins or partially unfolded hairpin conformations.

### IntI4 synthesis, expression and purification

IntI4 integrase coding sequence (Uniprot #O68847) with one nucleotide change (A904T) resulting in Tyr302Phe substitution was synthesized as gBlocks (IDT, USA) in fusion with a maltose binding protein (MBP) for solubility and stability purposes (full sequence in Table S4). The synthesized fragment was added to the pMAL-c5X plasmid using In-Fusion (Takara Bio, Japan) and the resulting plasmid was transformed into the *E. coli* BL21 (DE3) strain using heat-shock. Expression and purification process was identical to the one previously described in (Vorobevskaia et al. 2024) for MBP-IntI1 protein. In brief, a 1 L Terrific Broth (TB) culture containing 50 μg/mL ampicillin was inoculated with 5 mL of overnight culture and grown at 37°C, 140 rpm. At an optical density (OD_600_) of 0.6, protein expression was induced by adding 0.3 mM IPTG. The culture was incubated overnight at 14°C, then centrifuged at 5500 rpm for 30 min at 4°C. The pellet was resuspended, lysed using an EmulsiFlex-C3, and centrifuged at 8400 rpm for 1 hour at 4°C. The supernatant was filtered and loaded onto an ÄKTA FPLC system for MBPTrap affinity chromatography. The MBP-tagged mutant protein was eluted with a maltose-containing buffer, and fractions were analyzed by 12% SDS-PAGE. A second purification step using ion exchange chromatography (HiTrap SP HP, Cytiva, USA) removed remaining impurities. Fractions were confirmed by SDS-PAGE, dialyzed into storage buffer, snap-frozen, and stored at −70°C.

### Integron plasmid engineering

Based on a p15A integron plasmid (kind gift of Prof. Didier Mazel), we have modified the three last gene cassettes from the array substituting them with gene cassettes of our choice based on reported sequences of ORFs and accompanying *attC* sites in the INTEGRALL database (Moura et al. 2009). Briefly, the cassette array flanked with cloning overhangs was synthesized as a DNA fragment (2328 bp length; GenScript, USA) and inserted using the In-Fusion kit (Takara Bio, Japan). The resulting integron plasmid *p15A-IntI1wt-smR2-catB3-blaVIM4-aac(6’)-Ib-cr* (“*p15A-integron*”) was used in the experiments for strains #965 and #1008 (also #1008_IPTG_). The plasmid was introduced to the *E. coli MG1655* (both wild type and mCherry-pS-modified) using electroporation. As controls, we generated the same plasmid with an inactive integrase variant IntI1^Y312F^ using a QuikChange site-directed mutagenesis kit (Agilent, USA) on the *p15A-integron*. The primer design was done using QuikChange online tool (Table S3) and the reaction was carried out according to the manufacturer’s instructions. Mutations were confirmed using Sanger sequencing. The resulting plasmid *p15A-IntI1^Y312F^-smR2-catB3-blaVIM4-aac(6’)-Ib* was used in the strains #1009 (*MG1655_mCherry-pS_*) and #1010 (*MG1655_wt_*). We further generated a truncated integrase variant IntI1^ΔC^ plasmid using a Q5 site-directed mutagenesis kit (New England Biolabs, USA) on the *p15A-integron* by substituting Ala^321^ and Gly^322^ to stop codons (primer sequences in Table S3). The primer design was performed with NEBuilder online tool and the reaction was carried out according to the manufacturer’s instructions. Mutations were confirmed using Sanger sequencing. The resulting plasmid *p15A-IntI1 ^ΔC^-smR2-catB3-blaVIM4-aac(6’)-Ib-cr* – was used in the strain #1016 (*MG1655_wt_*).

### Introducing mCherry-pS through recombineering

The mCherry-pS construct was integrated into the *lacZ* locus of *E. coli* MG1655 using λ-Red recombineering as described earlier (Schärfen et al. 2020). Briefly, the auxiliary plasmid pR6K-mCherry-pS-lox71-cm-lox66 was assembled by In-Fusion cloning (Takara Bio, Japan). The *pR6K-lox71-cm-lox66* backbone was linearized by PCR using primer pR6K_vec_fwd and pR6K_vec_rev and the mCherry-pS insert was amplified from #956-*pMAL-5X_mCherry-pS* with 15-bp overlaps complementary to the vector ends using primer pS_mCherry_ins_fwd and pS_mCherry_ins_rev. The plasmid was assembled by In-Fusion reaction (15 min, 50 °C) and transformed into chemically competent *E. coli* DH5α-pir+ cells by heat shock. For chromosomal integration, the mCherry-pS-lox71-cm-lox66 cassette was PCR-amplified with primers mCherry_pS_rec_fwd and mCherry_pS_rec_rev carrying 50 bp homology region and 250 ng were electroporated into *E. coli* MG1655 harbouring *pSC101-BAD-γβαA*. Expression of λ-Red recombination genes (Redγβα) was induced with 0.25% L-arabinose prior to electroporation. Transformants were selected on LB agar supplemented with chloramphenicol (15 µg/ml) and screened by colony PCR to confirm correct insertion at the *lacZ* locus using primer lacZ_seq_fwd and lacZ_seq_rev. The chloramphenicol resistance cassette was excised with Cre-lox recombination using the temperature-sensitive plasmid *pSC101-BAD-Cre*. Cre expression was induced with 5 mM L-arabinose. Temperature-sensitive plasmids were removed during incubation at 37 °C.

### The in vivo adaptation assay under 1.3x MIC ciprofloxacin stress

The studied strains were grown in liquid culture in 1 mL volume of LB media supplemented with 50 µg/mL kanamycin (LB_Kana_) (resistance of the *p15A-integron* plasmid) and 1.3x MIC ciprofloxacin (40 ng/mL) to induce stress. The cultures were started using single colonies which were grown from the glycerol stock on Agar media supplemented with kanamycin. The liquid culture was grown for 19 hours (overnight) in aerated 1.5 mL Eppendorf tubes on a thermoshaker (Eppendorf, Germany) at 30°C with continuous orbital shaking at 500 rpm. The OD_600_ was measured for the overnight stressed cultures using a sterile 48 well plate (Greiner, Austria) in a CLARIOstar plate reader (BMG Labtech, Germany).

### Liquid culture PCR and agarose gel electrophoresis

The cultures were pelleted at 5000 g for 30 s and most supernatant was removed. The cells were re-suspended in the remaining media and 1 µL of the concentrated culture was used as a DNA template in a PCR reaction. The PCR mix consisted of the Quick-Load® Taq 2X Master Mix (New England Biolabs, USA), primers fwd_attI_EMSA and rev_aac6Ib-cr (Table S3) to amplify the region of interest from the cassette promoter Pcw to the end of the *aac(6’)-Ib-cr* gene cassette, 1 µL concentrated culture and milliQ. The reaction was carried out in 25 µL total volume and 6 µL of the PCR product were directly loaded on an agarose gel (0.8% in 1x TAE) pre-stained with ROTI GelStain (Carl Roth, Germany). The gel was visualized using a gel imager (Azure 300, Azure Biosystems, USA) and the size of the amplified bands was identified using the DNA ladder (GeneRuler DNA ladder Mix, ThermoFisher Scientific, USA).

### Dead cell assay

Dead cell assays were conducted on overnight cultures. Cells were pelleted at 5000 g for 30 s, the supernatant was discarded and the pellet re-suspended in 200 uL PBS. SYTOX Green stain was added to a final concentration of 1 µM and the staining proceeded in room temperature in the dark for 10 minutes. The cells were then washed with PBS three times using centrifugation. After the final wash the cells were re-suspended in 50 µL PBS and 1 µL of washed stained cultures was spread on an agar pad for imaging (Lörzing et al. 2024). Unstressed cells, grown without ciprofloxacin, were stain as control to determine the fraction of naturally occurring dead cells after overnight growth in experimental conditions and found between 1-2 % of dead cells. Additionally, a sample of the same unstressed culture was incubated at 75°C for 15 min for thermal killing to obtain a positive dead cell control leading to 100% dead cells. Dead cell assays were evaluated with fluorescence microscopy.

### Brightfield and Fluorescence Microscopy

Glass coverslips (№1.5) were cleaned by sonication twice for 15 min in 5 % (v/v) Mucasol solution, followed by a 15 min sonication in ethanol. After rinsing with milliQ water, coverslips were dried with nitrogen gas and stored dust free until use. Cells were immobilized on agarose pads as described in (Skinner et al. 2013). A 1 µl concentrated cell suspension was placed onto the agarose pad. The sample was covered with a cleaned coverslip and gently pressed to ensure uniform contact. Imaging was performed on a Nikon Eclipse Ti2-E inverted microscope equipped with an ORBITAL-100 Ring TIRF system (Visitron Systems, Germany), controlled using VisiView Software (Visitron Systems). Images were acquired with a 100x/ 1.49 NA Apo TIRF objective (Nikon) and an additional 1.5x tube lens (150x total magnification) and recorded with a Prime-95B Back-Illuminated scientific CMOS camera (Teledyne Photometrics, USA). For each field of view, a brightfield image was acquired prior to fluorescence imaging. Fluorescence imaging was performed using highly inclined and laminated optical sheet (HILO) illumination. mCherry was excited at 561 nm (8 mW, measured before the objective) and acquired as 50 frame time-lapse sequence at 50 frames per second. SYTOX Green was excited at 488 nm (0.2 mW, measured before the objective) and recorded as 50 frame time-lapse sequence at 20 frames per second. A quad-band filter set (ZET405/488/561/640m-TRFv2, Chroma) was used to filter emission light.

For mCherry analysis cells were segmented from brightfield images using a StarDist2D model trained with the ZeroCostDL4Mic platform (Lörzing et al. 2024; Schmidt et al. 2018; von Chamier et al. 2021). The resulting ROIs were applied to fluorescence images for quantification. Fifty frame time-lapse sequences were converted into average intensity projections in Fiji (ImageJ), and the mean fluorescence intensity within each ROI was calculated and used as the per-cell mCherry signal. Cell segmentation for stressed cells was performed manually in Fiji (ImageJ) using brightfield images. Individual cells were outlined with the ROI Manager, and the corresponding ROIs were applied to the fluorescence images. For SYTOX analysis, 50 frame time-lapse sequences were converted into average intensity projections, and fluorescence intensities were measured per cell. The maximum pixel intensity within each ROI was used for downstream quantification. Cell length was obtained from the “Major” axis parameter of the fitted ellipse. Subsequent data processing and analysis were performed using custom Python scripts. A SYTOX signal threshold was defined using the negative control (unstressed stained cells of #965 strain) as the mean maximum intensity plus two standard deviations. Cells with fluorescence intensities above this threshold were classified as SYTOX-positive. For each condition, the fraction of cells exceeding this threshold was calculated. Conversely, the fraction of cells below the threshold was defined as the surviving fraction.

## Author contributions

Conceptualization: MS, EV

Methodology: MS, PL

EV Investigation: EV

PL Visualization: EV, PL

Supervision: MS

Writing—original draft: EV

Writing—review & editing: EV, PL, MS

## Acknowledgments

We thank all members of the Schlierf lab for lively discussions during the development of this project. We thank Prof. Didier Mazel and Dr. Céline Loot (Institut Pasteur) for discussions and the initial integron plasmid, Prof. Francis Stewart and Dr. Frank Groß (TU Dresden) for the kind gift of the recombineering plasmids, Prof. Thorsten Mascher (TU Dresden) for the MG1655 strain, and Prof. Yixin Zhang (TU Dresden) for access to the plate reader. We acknowledge support by the PoL Microscopy facility, a core facility of the EXC PoL at TU Dresden. This research was funded by TU Dresden core funds (M.S.) and the BMFTR GO-Bio *initial* program (03LWH0050 to E.V.).

## Data and materials availability

All data are available in the main text or the supplementary materials. Raw data and analysis scripts are available upon request.

## Supplementary Materials

Supplementary Materials include Supplementary Text, Supplementary Figures, Supplementary Tables.

## References

Abramson, Josh, Jonas Adler, Jack Dunger, et al. 2024. “Accurate Structure Prediction of Biomolecular Interactions with AlphaFold 3.” Nature 630 (8016): 493–500. 10.1038/s41586-024-07487-w.

Aldred, Katie J., Robert J. Kerns, and Neil Osheroff. 2014. “Mechanism of Quinolone Action and Resistance.” Biochemistry 53 (10): 1565–74. 10.1021/bi5000564.

Ashkenazy, Haim, Shiran Abadi, Eric Martz, et al. 2016. “ConSurf 2016: An Improved Methodology to Estimate and Visualize Evolutionary Conservation in Macromolecules.” Nucleic Acids Research 44 (Web Server issue): W344–50. 10.1093/nar/gkw408.

Belan, Ondrej, George Moore, Artur Kaczmarczyk, et al. 2021. “Generation of Versatile Ss-dsDNA Hybrid Substrates for Single-Molecule Analysis.” STAR Protocols 2 (2): 100588. 10.1016/j.xpro.2021.100588.

Bos, Julia, Qiucen Zhang, Saurabh Vyawahare, Elizabeth Rogers, Susan M. Rosenberg, and Robert H. Austin. 2015. “Emergence of Antibiotic Resistance from Multinucleated Bacterial Filaments.” Proceedings of the National Academy of Sciences 112 (1): 178–83. 10.1073/pnas.1420702111.

Bouvier, Marie, Magaly Ducos-Galand, Céline Loot, David Bikard, and Didier Mazel. 2009. “Structural Features of Single-Stranded Integron Cassette attC Sites and Their Role in Strand Selection.” PLoS Genetics 5 (9): e1000632. 10.1371/journal.pgen.1000632.

Bustamante, Carlos, and Shannon Yan. 2022. “The Development of Single Molecule Force Spectroscopy: From Polymer Biophysics to Molecular Machines.” Quarterly Reviews of Biophysics 55: e9. 10.1017/S0033583522000087.

Cambray, Guillaume, Anne-Marie Guerout, and Didier Mazel. 2010. “Integrons.” Annual Review of Genetics 44 (1): 141–66. 10.1146/annurev-genet-102209-163504.

Castillo, Fabio, Amal Benmohamed, and George Szatmari. 2017. “Xer Site Specific Recombination: Double and Single Recombinase Systems.” Frontiers in Microbiology 8. https://www.frontiersin.org/articles/10.3389/fmicb.2017.00453.

Chamier, Lucas von, Romain F. Laine, Johanna Jukkala, et al. 2021. “Democratising Deep Learning for Microscopy with ZeroCostDL4Mic.” Nature Communications 12 (1): 2276. 10.1038/s41467-021-22518-0.

Chen, Ding-Qiang, Yue-Ting Jiang, Dong-Hua Feng, Shu-Xian Wen, Dan-Hong Su, and Ling Yang. 2018. “Integron Mediated Bacterial Resistance and Virulence on Clinical Pathogens.” Microbial Pathogenesis 114 (January): 453–57. 10.1016/j.micpath.2017.12.029.

Chen, Yu, Umadevi Narendra, Lisa E. Iype, Michael M. Cox, and Phoebe A. Rice. 2000. “Crystal Structure of a Flp Recombinase–Holliday Junction Complex: Assembly of an Active Oligomer by Helix Swapping.” Molecular Cell 6 (4): 885–97. 10.1016/S1097-2765(05)00088-2.

Collis, Christina M., and Ruth M. Hall. 1992. “Gene Cassettes from the Insert Region of Integrons Are Excised as Covalently Closed Circles.” Molecular Microbiology 6 (19): 2875–85. 10.1111/j.1365-2958.1992.tb01467.x.

Collis, Christina M., and Ruth M. Hall. 2004. “Comparison of the Structure–Activity Relationships of the Integron-Associated Recombination Sites attI3 and attI1 Reveals Common Features.” Microbiology 150 (5): 1591–601. 10.1099/mic.0.26596-0.

Darby, Elizabeth M., Eleftheria Trampari, Pauline Siasat, et al. 2023. “Molecular Mechanisms of Antibiotic Resistance Revisited.” Nature Reviews Microbiology 21 (5): 280–95. 10.1038/s41579-022-00820-y.

Darracq, Baptiste, Eloi Littner, Manon Brunie, et al. 2025. “Sedentary Chromosomal Integrons as Biobanks of Bacterial Antiphage Defense Systems.” Science 388 (6747): eads0768. 10.1126/science.ads0768.

Davies, Julian. 1994. “Inactivation of Antibiotics and the Dissemination of Resistance Genes.” Science 264 (5157): 375–82. 10.1126/science.8153624.

Escudero, José Antonio, Céline Loot, Aleksandra Nivina, and Didier Mazel. 2015. “The Integron: Adaptation On Demand.” Microbiology Spectrum 3 (2): 3.2.10. 10.1128/microbiolspec.MDNA3-0019-2014.

Gopaul, D. N., F. Guo, and G. D. Van Duyne. 1998. “Structure of the Holliday Junction Intermediate in Cre-loxP Site-Specific Recombination.” The EMBO Journal 17 (14): 4175–87. 10.1093/emboj/17.14.4175.

Guerin, Émilie, Guillaume Cambray, Neus Sanchez-Alberola, et al. 2009. “The SOS Response Controls Integron Recombination.” Science 324 (5930): 1034–1034. 10.1126/science.1172914.

Guérin, Emilie, Thomas Jové, Aurore Tabesse, Didier Mazel, and Marie-Cécile Ploy. 2011. “High-Level Gene Cassette Transcription Prevents Integrase Expression in Class 1 Integrons.” Journal of Bacteriology 193 (20): 5675–82. 10.1128/jb.05246-11.

Hansson, Karin, Ola Sköld, and Lars Sundström. 1997. “Non-palindromic *attI* Sites of Integrons Are Capable of Site-specific Recombination with One Another and with Secondary Targets.” Molecular Microbiology 26 (3): 441–53. 10.1046/j.1365-2958.1997.5401964.x.

Hickman, Alison Burgess, Shani Waninger, John J. Scocca, and Fred Dyda. 1997. “Molecular Organization in Site-Specific Recombination: The Catalytic Domain of Bacteriophage HP1 Integrase at 2.7 Å Resolution.” Cell 89 (2): 227–37. 10.1016/S0092-8674(00)80202-0.

Jayaram, Makkuni, Chien-Hui Ma, Aashiq H. Kachroo, et al. 2015. “An Overview of Tyrosine Site-Specific Recombination: From an Flp Perspective.” Microbiology Spectrum 3 (4): 10.1128/microbiolspec.mdna3-0021–2014. 10.1128/microbiolspec.mdna3-0021–2014.

Jové, Thomas, Sandra Da Re, François Denis, Didier Mazel, and Marie-Cécile Ploy. 2010. “Inverse Correlation between Promoter Strength and Excision Activity in Class 1 Integrons.” PLoS Genetics 6 (1). 10.1371/journal.pgen.1000793.

Kieffer, Nicolas, Alberto Hipólito, Laura Ortiz-Miravalles, et al. 2025. “Mobile Integrons Encode Phage Defense Systems.” Science 388 (6747): eads0915. 10.1126/science.ads0915.

Landy, Arthur. 2015. “The λ Integrase Site-Specific Recombination Pathway.” In Mobile DNA III. John Wiley & Sons, Ltd. 10.1128/9781555819217.ch4.

Loot, Céline, David Bikard, Anna Rachlin, and Didier Mazel. 2010. “Cellular Pathways Controlling Integron Cassette Site Folding.” The EMBO Journal 29 (15): 2623–34. 10.1038/emboj.2010.151.

Loot, Céline, Gael A. Millot, Egill Richard, et al. 2024. “Integron Cassettes Integrate into Bacterial Genomes via Widespread Non-Classical attG Sites.” Nature Microbiology 9 (1): 228–40. 10.1038/s41564-023-01548-y.

Lörzing, Pilar, Philipp Schake, and Michael Schlierf. 2024. “Anisotropic DBSCAN for 3D SMLM Data Clustering.” The Journal of Physical Chemistry B 128 (33): 7934–40. 10.1021/acs.jpcb.4c02030.

MacDonald, Douglas, Gaëlle Demarre, Marie Bouvier, Didier Mazel, and Deshmukh N. Gopaul. 2006. “Structural Basis for Broad DNA-Specificity in Integron Recombination.” Nature 440 (7088): 1157–62. 10.1038/nature04643.

Madeira, Fábio, Nandana Madhusoodanan, Joonheung Lee, et al. 2024. “The EMBL-EBI Job Dispatcher Sequence Analysis Tools Framework in 2024.” Nucleic Acids Research 52 (W1): W521–25. 10.1093/nar/gkae241.

Mazel, Didier. 2006. “Integrons: Agents of Bacterial Evolution.” Nature Reviews Microbiology 4 (8): 608–20. 10.1038/nrmicro1462.

Mazel, Didier, Broderick Dychinco, Vera A. Webb, and Julian Davies. 1998. “A Distinctive Class of Integron in the Vibrio Cholerae Genome.” Report. Science 280 (5363): 605–8. 10.1126/science.280.5363.605.

Meinke, Gretchen, Andrew Bohm, Joachim Hauber, M. Teresa Pisabarro, and Frank Buchholz. 2016. “Cre Recombinase and Other Tyrosine Recombinases.” Chemical Reviews 116 (20): 12785–820. 10.1021/acs.chemrev.6b00077.

Michel, Bénédicte. 2005. “After 30 Years of Study, the Bacterial SOS Response Still Surprises Us.” PLoS Biology 3 (7). 10.1371/journal.pbio.0030255.

Moura, Alexandra, Mário Soares, Carolina Pereira, Nuno Leitão, Isabel Henriques, and António Correia. 2009. “INTEGRALL: A Database and Search Engine for Integrons, Integrases and Gene Cassettes.” Bioinformatics (Oxford, England) 25 (8): 1096–98. 10.1093/bioinformatics/btp105.

Mukhortava, Ann, Matthias Pöge, Maj Svea Grieb, et al. 2019. “Structural Heterogeneity of *attC* Integron Recombination Sites Revealed by Optical Tweezers.” Nucleic Acids Research 47 (4): 1861–70. 10.1093/nar/gky1258.

Murray, Christopher J. L., Kevin Shunji Ikuta, Fablina Sharara, et al. 2022. “Global Burden of Bacterial Antimicrobial Resistance in 2019: A Systematic Analysis.” The Lancet 399 (10325): 629–55. 10.1016/S0140-6736(21)02724-0.

Naghavi, Mohsen, Stein Emil Vollset, Kevin S. Ikuta, et al. 2024. “Global Burden of Bacterial Antimicrobial Resistance 1990–2021: A Systematic Analysis with Forecasts to 2050.” The Lancet 404 (10459): 1199–226. 10.1016/S0140-6736(24)01867-1.

Nandi, Sobhan, John J. Maurer, Charles Hofacre, and Anne O. Summers. 2004. “Gram-Positive Bacteria Are a Major Reservoir of Class 1 Antibiotic Resistance Integrons in Poultry Litter.” Proceedings of the National Academy of Sciences of the United States of America 101 (18): 7118–22. 10.1073/pnas.0306466101.

Nivina, Aleksandra, José Antonio Escudero, Claire Vit, Didier Mazel, and Céline Loot. 2016. “Efficiency of Integron Cassette Insertion in Correct Orientation Is Ensured by the Interplay of the Three Unpaired Features of attC Recombination Sites.” Nucleic Acids Research 44 (16): 7792–803. 10.1093/nar/gkw646.

Partridge, Sally R., Gavin D. Recchia, Carol Scaramuzzi, Christina M. Collis, H. W. Stokes, and Ruth M. Hall. 2000. “Definition of the attI1 Site of Class 1 Integrons.” Microbiology 146 (11): 2855–64. 10.1099/00221287-146-11-2855.

Phillips, Ian, Esther Culebras, Felipe Moreno, and Fernando Baquero. 1987a. “Induction of the SOS Response by New 4-Quinolones.” Journal of Antimicrobial Chemotherapy 20 (5): 631–38. 10.1093/jac/20.5.631.

Richard, Egill, Baptiste Darracq, Céline Loot, and Didier Mazel. 2022. “Unbridled Integrons: A Matter of Host Factors.” Cells 11 (6): 925. 10.3390/cells11060925.

Rowe-Magnus, Dean A., Anne-Marie Guerout, and Didier Mazel. 2002. “Bacterial Resistance Evolution by Recruitment of Super-Integron Gene Cassettes.” Molecular Microbiology 43 (6): 1657–69. 10.1046/j.1365-2958.2002.02861.x.

Sallen, B., A. Rajoharison, S. Desvarenne, and C. Mabilat. 1995. “Molecular Epidemiology of Integron-Associated Antibiotic Resistance Genes in Clinical Isolates of Enterobacteriaceae.” Microbial Drug Resistance (Larchmont, N.Y.) 1 (3): 195–202. 10.1089/mdr.1995.1.195.

Santajit, Sirijan, and Nitaya Indrawattana. 2016. “Mechanisms of Antimicrobial Resistance in ESKAPE Pathogens.” BioMed Research International 2016 (1): 2475067. 10.1155/2016/2475067.

Schärfen, Leonard, Miloš Tišma, and Michael Schlierf. 2020. “Fast, Simultaneous Tagging and Mutagenesis of Genes on Bacterial Chromosomes.” ACS Synthetic Biology 9 (8): 2203–7. 10.1021/acssynbio.0c00202.

Schmidt, Uwe, Martin Weigert, Coleman Broaddus, and Gene Myers. 2018. “Cell Detection with Star-Convex Polygons.” In Medical Image Computing and Computer Assisted Intervention – MICCAI 2018, edited by Alejandro F. Frangi, Julia A. Schnabel, Christos Davatzikos, Carlos Alberola-López, and Gabor Fichtinger. Springer International Publishing. 10.1007/978-3-030-00934-2_30.

Schrödinger, L., & DeLano, W. 2020. PyMOL. Released. http://www.pymol.org/pymol.

Skinner, Samuel O., Leonardo A. Sepúlveda, Heng Xu, and Ido Golding. 2013. “Measuring mRNA Copy Number in Individual Escherichia Coli Cells Using Single-Molecule Fluorescent in Situ Hybridization.” Nature Protocols 8 (6): 1100–1113. 10.1038/nprot.2013.066.

Spåhr, Henrik, Tiongsun Chia, James P. Lingford, et al. 2018. “Modular ssDNA Binding and Inhibition of Telomerase Activity by Designer PPR Proteins.” Nature Communications 9 (1): 1. 10.1038/s41467-018-04388-1.

Stokes, H. W., and R. M. Hall. 1989. “A Novel Family of Potentially Mobile DNA Elements Encoding Site-Specific Gene-Integration Functions: Integrons.” Molecular Microbiology 3 (12): 1669–83. 10.1111/j.1365-2958.1989.tb00153.x.

Subramanya, Hosahalli S., Lidia K. Arciszewska, Rachel A. Baker, Louise E. Bird, David J. Sherratt, and Dale B. Wigley. 1997. “Crystal Structure of the Site-Specific Recombinase, XerD.” The EMBO Journal 16 (17): 5178–87. 10.1093/emboj/16.17.5178.

Swoboda, Marko, Jörg Henig, Hsin-Mei Cheng, et al. 2012. “Enzymatic Oxygen Scavenging for Photostability without pH Drop in Single-Molecule Experiments.” ACS Nano 6 (7): 6364–69. 10.1021/nn301895c.

Taylor, Arnulf M. K., Stephen R. Okoniewski, Lyle Uyetake, and Thomas T. Perkins. 2022. “Force-Activated DNA Substrates for In Situ Generation of ssDNA and Designed ssDNA/dsDNA Structures in an Optical-Trapping Assay.” In Optical Tweezers: Methods and Protocols, edited by Arne Gennerich. Springer US. 10.1007/978-1-0716-2229-2_10.

Teichmann, Lisa, Sam Luitwieler, Johan Bengtsson-Palme, and Benno ter Kuile. 2025. “Fluoroquinolone-Specific Resistance Trajectories in E. Coli and Their Dependence on the SOS-Response.” BMC Microbiology 25 (1): 37. 10.1186/s12866-025-03771-5.

Vikedal, Krister, Synnøve Brandt Ræder, Ida Mathilde Marstein Riisnæs, et al. 2025. “RecN and RecA Orchestrate an Ordered DNA Supercompaction Response Following Ciprofloxacin-Induced DNA Damage in Escherichia Coli.” Nucleic Acids Research 53 (10): gkaf437. 10.1093/nar/gkaf437.

Vorobevskaia, Ekaterina, Céline Loot, Didier Mazel, and Michael Schlierf. 2024. “The Recombination Efficiency of the Bacterial Integron Depends on the Mechanical Stability of the Synaptic Complex.” Science Advances 10 (50): eadp8756. 10.1126/sciadv.adp8756.

Wang, M. D., H. Yin, R. Landick, J. Gelles, and S. M. Block. 1997. “Stretching DNA with Optical Tweezers.” Biophysical Journal 72 (3): 1335–46. 10.1016/S0006-3495(97)78780-0.

World Health Organization. 2014. Antimicrobial Resistance: Global Report on Surveillance. World Health Organization. https://apps.who.int/iris/handle/10665/112642.

