## Supplementary Information for "Active destabilization of the integron synaptic complex reduces bacterial adaptation to antibiotics"

### Supplementary Text

*DMSO 1% control for peptide OT experiments*

Peptides were obtained commercially (Genscript, USA) and shipped dry. They were dissolved in DMSO to a concentration of 1 mM, which was further diluted 1:100 (or prediluted for titration experiments) for the optical tweezers measurements, resulting in 1% DMSO in the final peptide-protein sample. To investigate the possible influence of 1% DMSO on integrase, its synaptic complex assembly and stability, we performed additional control experiments without the peptide, but in presence of 1% DMSO in the integrase solution. We gathered a control dataset of comparable size to our reference IntI1-$attC_{aadA7}^{bs}$ synaptic complex (171 disassembly traces versus 169 events, respectively). The disassembly was $\overline{F}_{\text{diss}}$ (IntI1-$attC_{aadA7}^{bs}$ w. 1% DMSO) = 14.6 pN ± 0.7 pN (mean ± SEM) and according to the Welch T-test (Freedman et al. 2007) and the Kolmogorov-Smirnov test no significant difference was detected. We conclude that the presence of 1% DMSO during the measurements had no significant effect on the mechanical stability of synaptic complexes.

### Supplementary Figures:

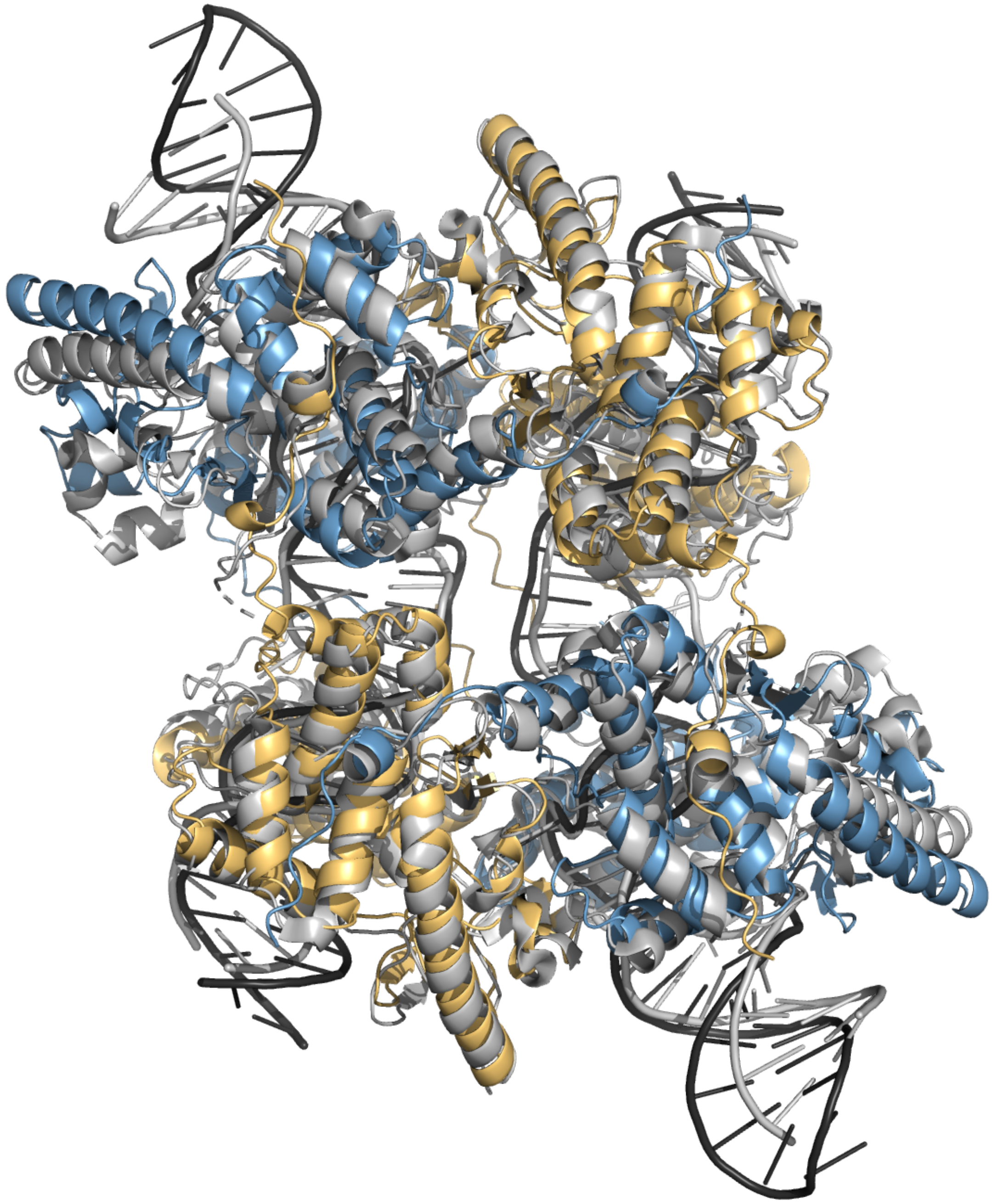

**Figure S1.** The AlphaFold3 predicted structure of the IntI1-*attC*$_{aadA7}$$^{bs}$ synaptic complex (Abramson et al. 2024), IntI1 monomers are indicated in orange and blue, DNA is black; aligned to the experimentally resolved crystallographic structure of the synaptic complex for the homologous integrase – IntI4 (MacDonald et al. 2006), colored grey. The alignment between both structures has the RMSD of 2.5 Å. Created using pyMOL (Schrödinger, L., & DeLano, W. 2020).

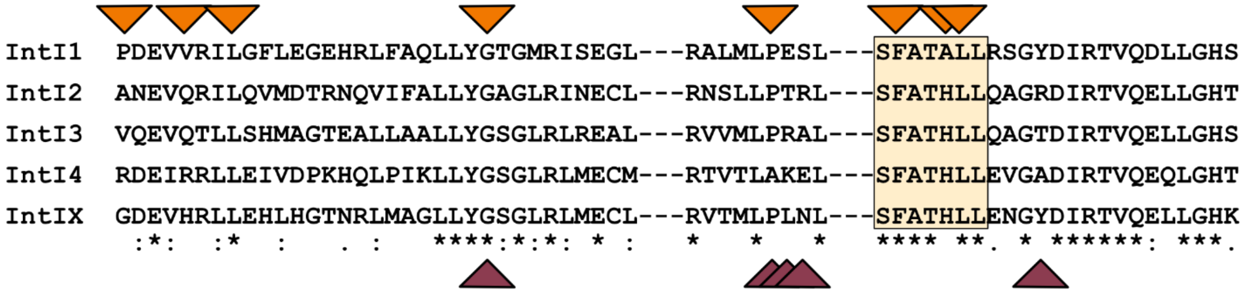

**Figure S2. Binding pocket alignment between different IntI classes**. Conserved residues are indicated with a star (*), red triangles indicate residues that form polar interactions, orange triangle – that form hydrophobic contact to the C-terminus tail. The docking dip for the $Pro^{326}_{Cterm}$ is highlighted in yellow. Binding pocket sequences were aligned using Clustal Omega Multiple Sequence Alignment (MSA) (Madeira et al. 2024). There is 50-60% sequence similarity between the integrase classes.

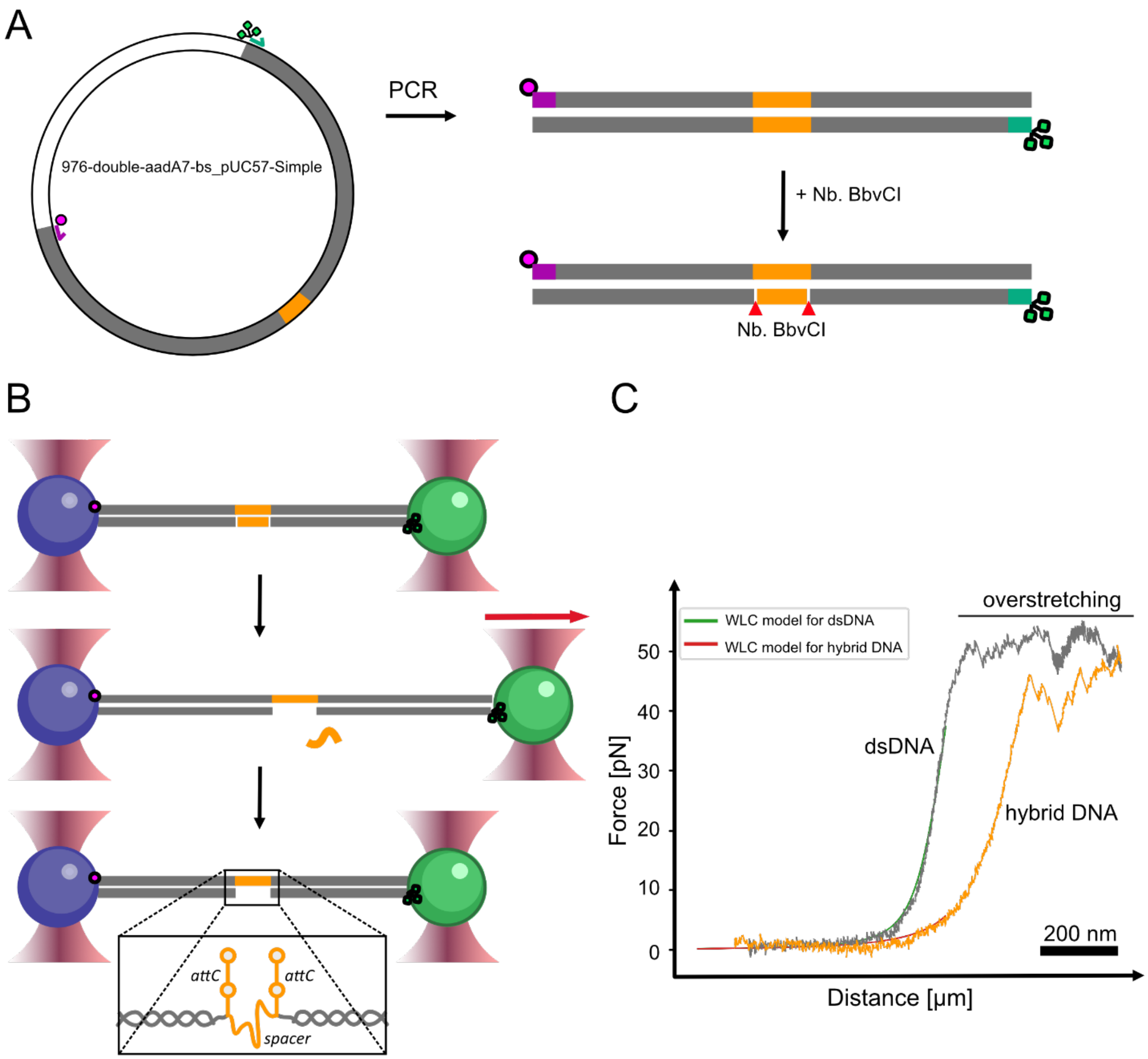

**Figure S3. Scheme of the hybrid DNA tether generation. (A)** dsDNA nicked construct preparation. The dsDNA construct is amplified from the plasmid using biotin and triple-digoxigenin functionalized primers, indicated in violet and green. The construct is nicked using the Nb. BbvCI enzyme and then cleaned using a kit. **(B)** The dsDNA nicked construct is tethered between the functionalized beads trapped by the optical tweezers and the DNA is overstretched by pulling one trap away to peel off the DNA strand between the nicks. The hybrid DNA is used for the synaptic reconstitution with the ssDNA presenting two $attC_{aadA7}{}^{bs}$ hairpins for IntI binding and a spacer for gene cassette imitation. **(C)** Exemplary force-extension curve showing the transition from dsDNA-like behavior (grey) to hybrid dsDNA-ssDNA behavior (orange).

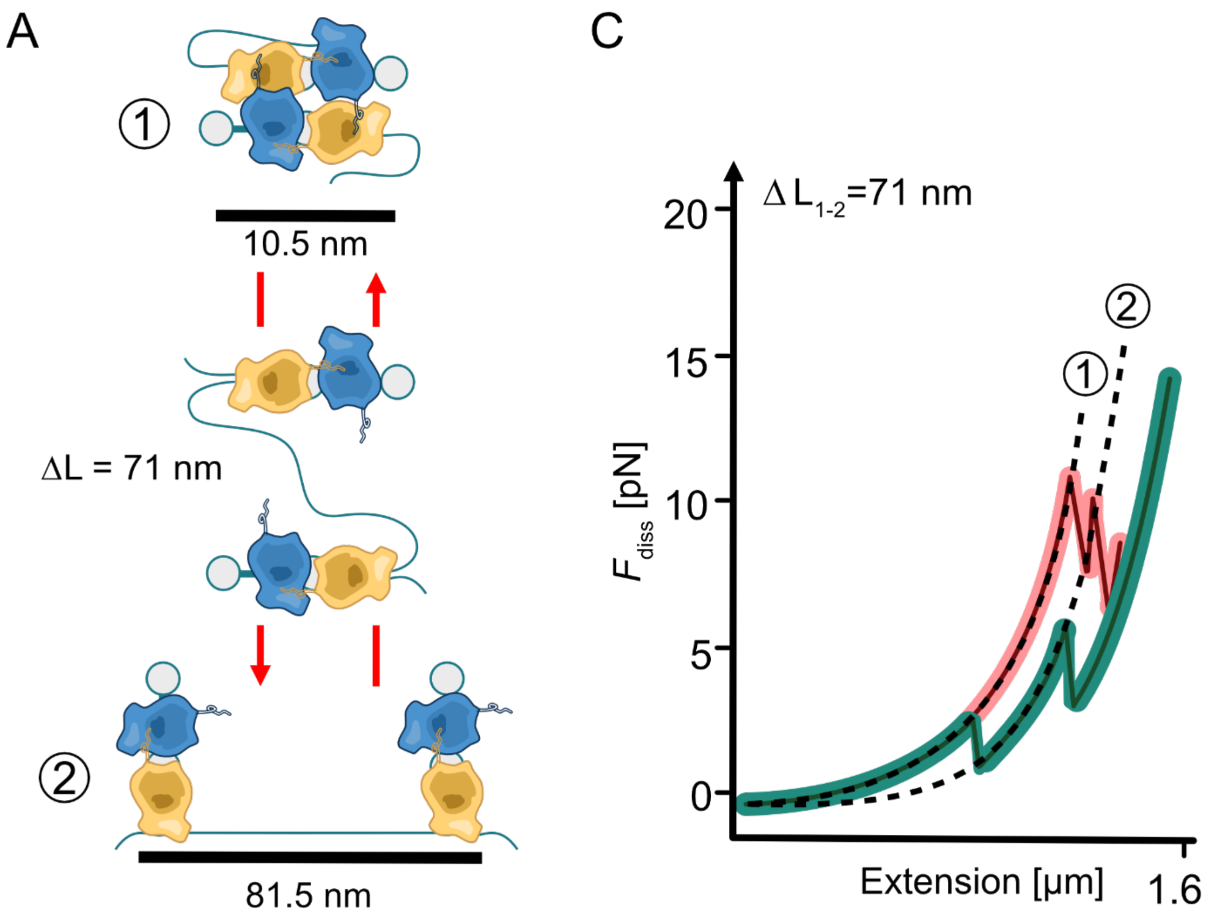

**Figure S4. Characteristic molecular fingerprint of the synaptic complex disassembly. (A)** The synaptic complex disassembly scheme from the formed complex (1) to the next stable state – both *attC* hairpins are bound and the spacer is stretched (2). The calculated characteristic contour length change $\Delta L_{1\text{-}2}$=71 nm. **(B)** Characteristic force-extension curves for high (in red) and low (in teal) stability synaptic complexes. The synapse disassembly is identified by the characteristic molecular fingerprint $\Delta L_{1\text{-}2}$=71 nm, and the disassembly force is recorded using the breaking point of the state (1) on the force-extension curve.

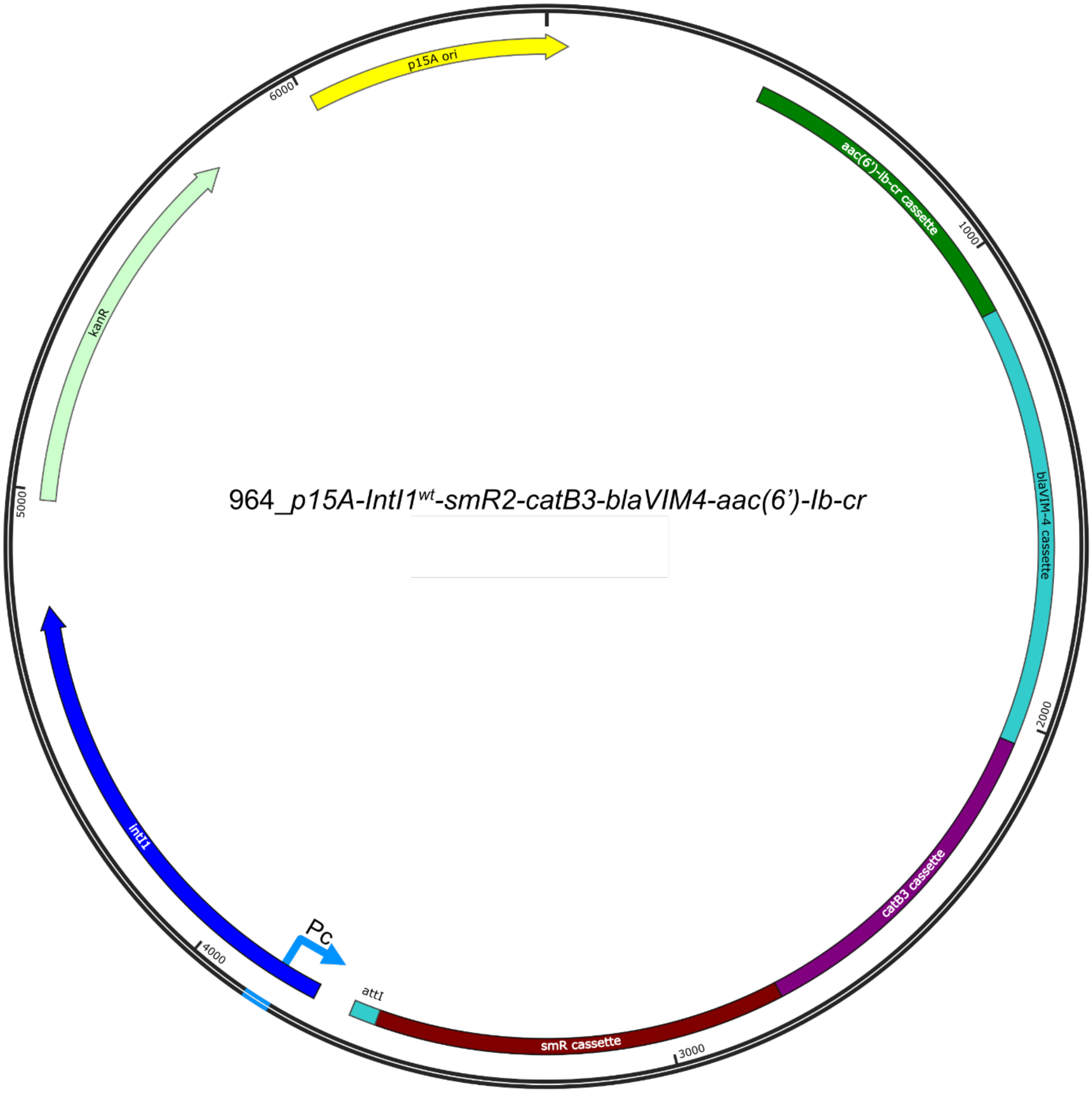

**Figure S5. The plasmid map of the p15A-integron plasmid.** The plasmid carries a *kanR* (kanamycin) selectable marker and a p15A origin of replication. The class 1 integron has the intI1 gene, *attI* site and a library of promoterless gene cassettes that include an ORF for an antibiotic resistance as well as the corresponding *attC* site for recombination. The cassettes are located in the following order from the Pc promoter region: *smR* – resistance against streptomycin; *catB3* – resistance against chloramphenicol; *blaVIM-4* – resistance against carbopenems and other beta-lactams; *aac(6')-Ib-cr* – resistance against ciprofloxacin and norfloxacin. Created with SnapGene Viewer (SnapGene, USA).

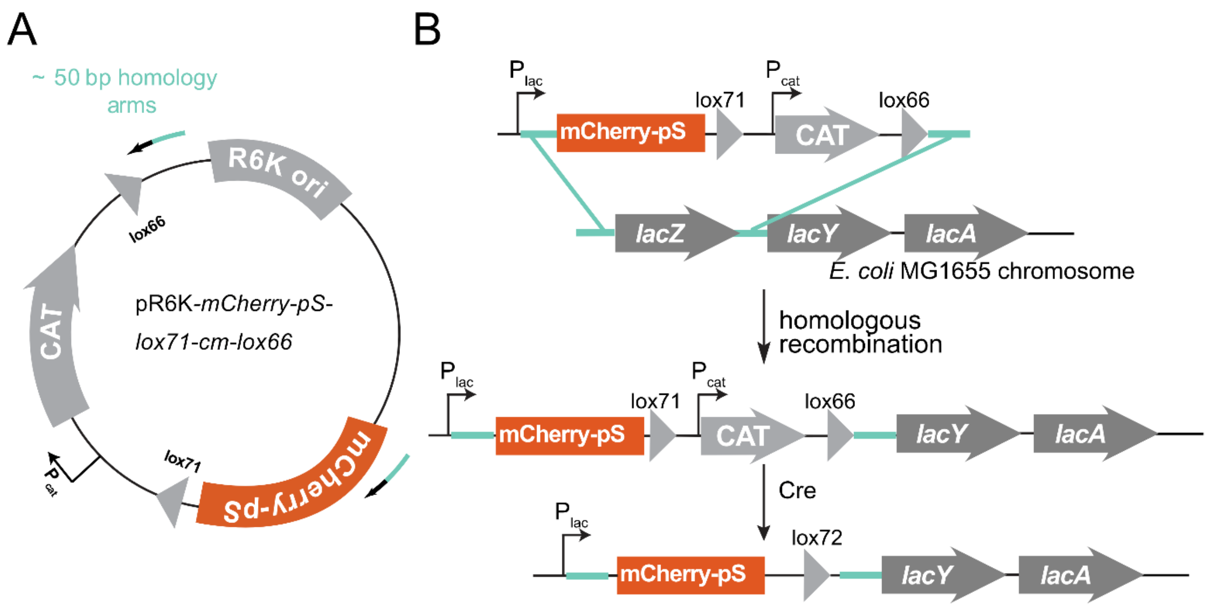

**Figure S6. mCherry-pS insertion on the *E.coli* MG1655 chromosome using Lambda Red recombineering. (A)** The targeting vector was constructed by cloning the mCherry-pS sequence (orange) into a pR6K backbone containing a chloramphenicol resistance gene (*cat*). The vector is then linearized by PCR, using primers that added 50-bp homology regions (turquoise) matching the target site in the genome for targeted genomic insertion. **(B)** A targeting vector is introduced into the cell. The Lambda Red system mediates homologous recombination to insert the mCherry-pS sequence replacing the *lacZ* gene in the lac operon. The chloramphenicol cassette is subsequently removed via Cre–loxP recombination.

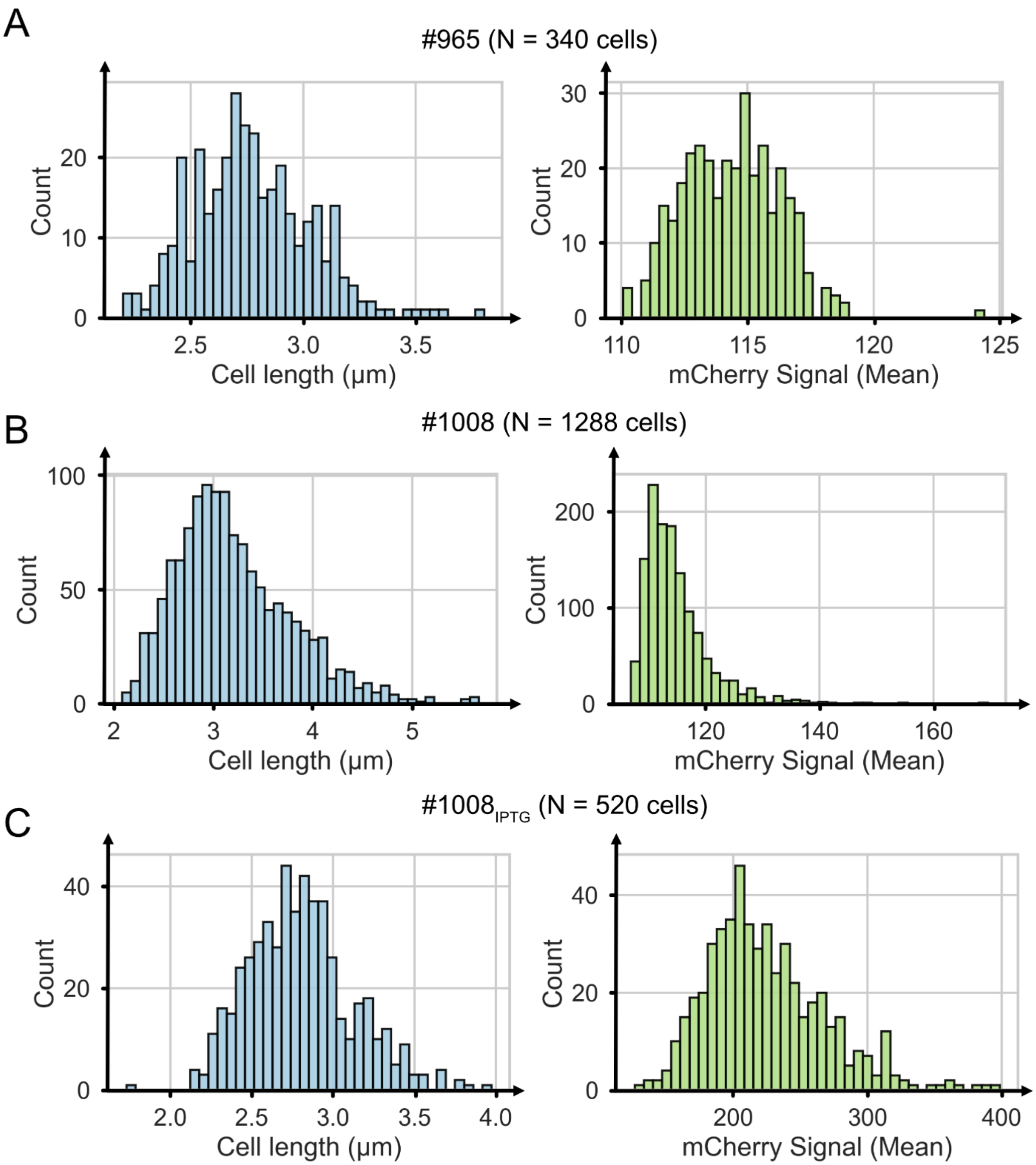

**Figure S7. The histograms of cell length and raw intensity of mCherry signal for the unstressed cells.** Bacterial strains and the number of analyzed cells are indicated, the mCherry signal is reported in arbitrary units as a mean value per cell.

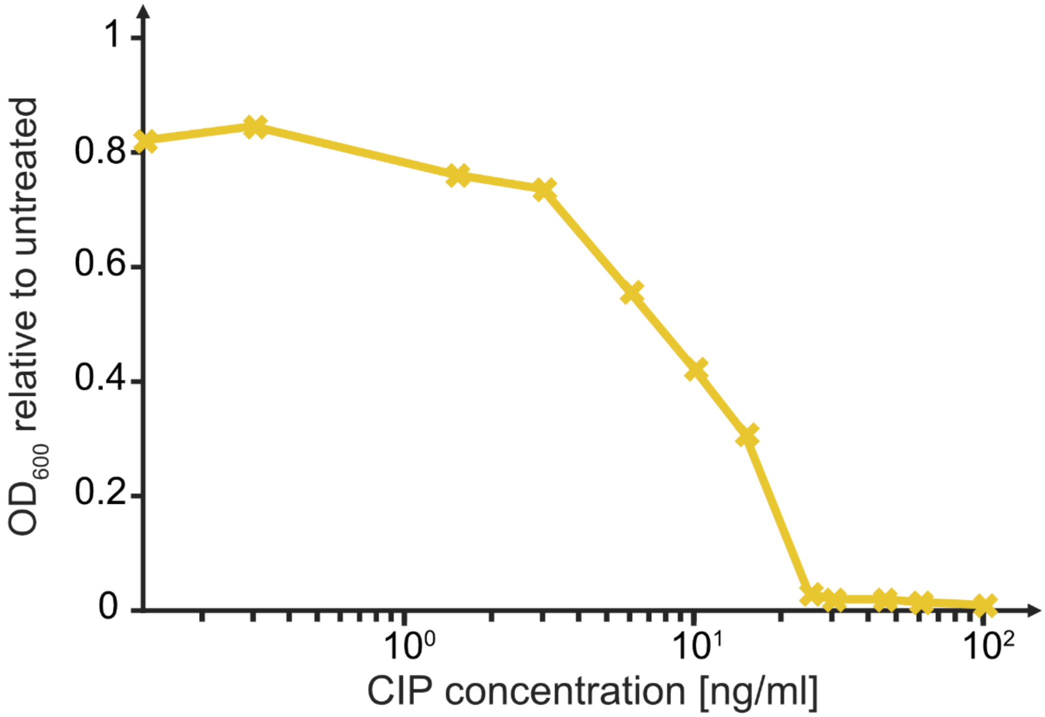

**Figure S8. The ciprofloxacin MIC determination for *E. coli* MG1655 using serial dilutions in LB medium.** Ciprofloxacin was twofold serially diluted in LB medium, inoculated with pre-grown *E. coli* culture and incubated o/n under standard growth conditions. MIC was defined as the lowest concentration at which no visible bacterial growth was observed – 32 ng/mL.

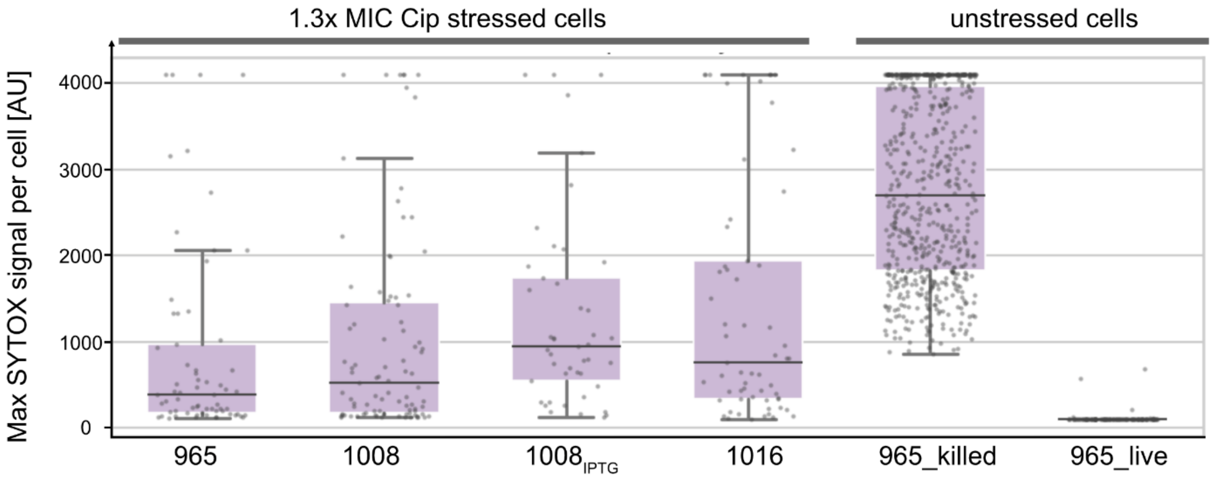

**Figure S9. The box plots of raw intensity of SYTOX signal.** Both stressed (1.3x MIC ciprofloxacin) and unstressed cells are shown. The unstressed cells from strain #965 are used as controls: heat-killed cells represent maximum SYTOX staining levels and live cells show background signal of membrane-intact cells. The SYTOX signal is reported in arbitrary units as a maximum value per cell.

### Supplementary Tables

**Table S1. Measurement data summary of the single-molecule optical tweezers experiments.** All synaptic complexes were formed using the *aadA7*$^{bs}$ *attC* site. Calculated median, mean force, and the SEM (standard error of the mean) are presented for all synaptic complex disassembly events.

| Protein | + peptide (concentration) | # Synaptic complex disassembly | Median force (pN) | Mean force (pN) | SEM (pN) |
|---|---|---|---|---|---|
| IntI1 | - | 169 | 12.7 | 13.9 | 0.6 |
| | pL (10 μM) | 178 | 8.9 | 10.2 | 0.5 |
| | pS (2 μM) | 164 | 12.6 | 15.0 | 0.7 |
| | pS (5 μM) | 106 | 9.9 | 11.7 | 0.6 |
| | pS (10 μM) | 221 | 9.2 | 10.4 | 0.4 |
| | pS (50 μM) | 113 | 9.1 | 10.1 | 0.5 |
| | pS’ (10 μM) | 108 | 11.5 | 12.7 | 0.6 |
| | pS’’ (10 μM) | 114 | 8.3 | 9.2 | 0.4 |
| | pS4 (10 μM) | 103 | 9.4 | 10.9 | 0.6 |
| IntI4 | - | 234 | 15.0 | 16.4 | 0.6 |
| | pS (10 μM) | 203 | 12.6 | 12.7 | 0.4 |

**Table S2. Data summary for microbiology experiments. (A)** Characteristics of unstressed cells of the studied strains. Integrase is always the wild type ($IntI1^{wt}$), $OD_{600}$, growth rate (r, calculated using Richards growth model (Zwietering et al. 1990)) and the cell length are given as mean ± SD. The raw mCherry signal is given in arbitrary units, mean ± SEM. Survival fractions were calculated for strain #965 in two conditions: live cells stained and heat-killed cells stained with SYTOX Green dye. **(B)** Characteristics of the 1.3x MIC ciprofloxacin stressed cells of the studied strains. Integrase variants are indicated, $OD_{600}$ and the cell length are given as mean ± SEM. The raw SYTOX Green signal is given in arbitrary units, mean ± SEM. Survival fractions were calculated for all strains by direct staining with SYTOX Green dye.

**A.** Characteristics of the unstressed cells of the studied strains

| Strain | mCherry-pS | $OD_{600}$ | Growth rate (r) | Cell length [µm] | Raw mCherry signal [AU] | Survival fraction live/killed |
|---|---|---|---|---|---|---|
| #965 | - | 2.55 ± 0.25 | 0.33 ± 0.01 | 2.8 ± 0.3 | 114.4 ± 1.9 | 0.98/0.00 |
| #1008 | + | 2.47 ± 0.21 | 0.29 ± 0.01 | 3.2 ± 0.6 | 114.7 ± 5.8 | - |
| $\#1008_{IPTG}$ | + | 2.59 ± 0.22 | 0.31 ± 0.01 | 2.8 ± 0.3 | 224.1 ± 43.9 | - |

**B.** Characteristics of the 1.3x MIC ciprofloxacin stressed cells of the studied strains

| Strain | mCherry-pS | IntI1 | $OD_{600}$ | Cell length [µm] | Raw SYTOX signal [AU] | Survival fraction |
|---|---|---|---|---|---|---|
| #965 | - | $IntI1^{wt}$ | 0.54 ± 0.05 | 13.1 ± 0.8 | 864.6 ± 138.4 | 0.4 |
| #1008 | + | $IntI1^{wt}$ | 0.49 ± 0.05 | 14.6 ± 0.8 | 1025.6 ± 123.7 | 0.34 |
| $\#1008_{IPTG}$ | + | $IntI1^{wt}$ | 0.37 ± 0.04 | 17.1 ± 1.1 | 1295.5 ± 167.9 | 0.11 |
| #1016 | - | $IntI1^{\Delta C}$ | 0.06 ± 0.03 | 21.3 ± 1.0 | 1410.8 ± 179.4 | 0.20 |

**Table S3. DNA sequences used in this study. (A)** Primers for dsDNA double-$attC_{aadA7}^{bs}$ construct. **(B)** Primers for cloning, mutagenesis, recombineering and liquid culture PCR. **(C)** Novel design of the double-$attC_{aadA7}^{bs}$ insert sequence flanked by Nb. BbvCI nicking sites. **(D)** Plasmids used for cloning, mutagenesis and recombineering. **(E)** Bacterial strains used in this study. **(F)** Constructed strains for *in vivo* adaptation assay.

**A.** Primers designed for PCR amplification of dsDNA double-$attC_{aadA7}^{bs}$ construct. Sequences are given in a 5' to 3' direction.

| **Biotin and triple-digoxigenin functionalized dsDNA construct** | |
|---|---|
| biotin_fwd_852bp_handle_AM | [Biotin] - CAGCATTGGTGACCTTGTTC |
| Triple-dig_fwd | [Triple-DIG] - ATCCGCAGAAGACGCAGATGCC |

**B.** Primers used in this study. Sequences are given in a 5' to 3' direction.

| **Plasmid p15A linearization** | |
|---|---|
| for_vector_930 | TTAGGTATGGATCCCATGGTACGC |
| rev_vector_930 | CGCTTGAGTTAAGCCGCG |
| **Protein mutagenesis ($IntI1^{Y312F}$ and $IntI1^{\Delta C}$)** | |
| L226_mut_IntI1 | GACGTCTCTACGACGATGATTTTCACGCATGTGCT |
| primer_fwd_mut_stop | GTGCTGAAAGTTGGCGGTTAGTGAGTGCGCTCACCGCTTG |
| **Primers used for pR6K-mCherry-pS cloning** | |
| pR6K_vec_fwd | CTGCTAAAGGAAGCGTACCGT |
| pR6K_vec_rev | CCCTTGCGCCCTGAGTG |
| pS_mCherry_ins_fwd | CTCAGGGCGCAAGGGATGGTGTCAAAGGGAGAGGAGGATAACATGG |
| pS_mCherry_ins_rev | CGCTTCCTTTAGCAGCTACAGCGCATCAAGCGGT |
| **Primers used for recombineering; 50 bp homology region is underlined** | |
| mCherry-pS-rec_fwd | TATGTTGTGTGGAATTGTGAGCGGATAACAATTTCACACAGGAAACAGCTATGGTGTCAAAGGGAGAGGAGG |
| mCherry-pS-rec_rev | ATGGATTTCCTTACGCGAAATACGGGCAGACATGGCCTGCCCGGTTATTAGCAGGATAGGTGAAGTAGGTACCG |
| **Primers used for colony PCR of lac operon** | |
| lacZ_seq_fwd | CACCCCAGGCTTTACACTT |
| lacZ_seq_rev | CAAATCGGGAAAAACGGGAAG |
| **Liquid culture PCR primers** | |
| fwd_attI_EMSA | GTTACGCCGTGGGTCG |
| rev_aac6Ib-cr | TTAGGCATCACTGCGTGTTC |

**C.** Novel design of the double-*attC*$_{aadA7}^{bs}$ insert sequence flanked by Nb. BbvCI nicking sites. The sequence of an *attC* site is indicated in red; Nb. BbvCI nicking sites are in green. The nick happens on the second strand between T and CG indicated by in the sequence by “|”. Sequences are given in a 5’ to 3’ direction.

<table>
<tr><td>Double-attC$_{aadA7}^{bs}$</td><td>CCTCA|GCATCTTTTATGTCTAACGCTTGAATTAAGCCGCGCCGCGAAGCGGCGTCGGCTTGAATGA<br>ATTGTTAGACATTTTCCATGGCTATATTTCAGCATCATCACATCATCATCATCATCATCACAGGGTA<br>GATCATCATCATCAACACGGGACGACAGCAAATGGCACCCTTCTATAAGCTTTTTATGTCTAACGCT<br>TGAATTAAGCCGCGCCGCGAAGCGGCGTCGGCTTGAATGAATTGTTAGACATTTTCCCTCA|GC</td></tr>
</table>

**D.** Plasmids used in this study.

| **Plasmid name** | **Description** | **Reference** |
|---|---|---|
| **Plasmids for double-*attC*$_{aadA7}^{bs}$ construct** | | |
| 3Dig-aadA7-bs_pUC57-Simple | Containing one handle and *attC*$_{aadA7}^{bs}$ | This study |
| 976-double-aadA7-bs_pUC57-Simple | Containing full double-*attC*$_{aadA7}^{bs}$ construct | This study |
| **Plasmids for integrase expression** | | |
| pMAL_c5X_IntI_Y312F_MBP | Catalytically inactive MBP-IntI1$^{Y312F}$ expression plasmid | (Grieb et al. 2017) |
| pMAL_c5X_IntI4_Y302F_MBP | Catalytically inactive MBP-IntI4$^{Y302F}$ expression plasmid | This study |
| **Plasmids for recombineering** | | |
| 956-pMAL-5X_mCherry-pS | Containing mCherry-pS | This study |
| pR6K-lox71-cm-lox66 | Target vector for recombineering | Kind gift from Prof. A. Francis Stewart |
| pR6K-mCherry-pS-lox71-cm-lox66 | Auxiliary plasmid for recombineering | This study |
| pSC101-BAD-γβαA | Expression plasmid for Red recombineering system | Kind gift from Prof. A. Francis Stewart |
| pSC101-BAD-Cre | Expression plasmid for Cre recombinase | Kind gift from Prof. A. Francis Stewart |
| **Plasmids for adaptation assay** | | |
| p15A-original integron | Carrying class 1 integron with four resistance gene cassettes in the library | Kind gift from Prof. Didier Mazel |

| | | |
|---|---|---|
| 964_p15A-IntI1$^{wt}$-smR2-catB3-blaVIM4-aac(6')-Ib-cr | Carrying class 1 integron with three new gene cassettes (*catB3-blaVIM4-aac(6')-Ib-cr*) | This study |
| 977_p15A-IntI1$^{Y312F}$-smR2-catB3-blaVIM4-aac(6')-Ib-cr | Carrying class 1 integron with three new gene cassettes and mutated inactive integrase IntI1$^{Y312F}$ | This study |
| 1015_p15A-IntI1$^{\Delta C}$-smR2-catB3-blaVIM4-aac(6')-Ib-cr | Carrying class 1 integron with three new gene cassettes and truncated integrase IntI1$^{\Delta C}$ | This study |

**E.** Bacterial strains used in this study.

| **Bacterial strain** | **Description** | **Reference** |
|---|---|---|
| *E. coli* MG1655 | F- λ- rph-1 | Kind gift from Prof. Thorsten Mascher |
| *E. coli* MG1655 ΔlacZ::mCherry-pS-lox71-cm-lox66 | F- λ- rph-1 *ΔlacZ::mCherry-pS-lox71-cm-lox66* | This study |
| *E. coli* MG1655 ΔlacZ::mCherry-pS-lox71/66 | F- λ- rph-1 *ΔlacZ::mCherry-pS-lox71/66* | This study |
| *E. coli* DH5α-pir$^{+}$ | *endA1 hsdR17 glnV44 (= supE44) thi-1 recA1 gyrA96 relA1 φ80dlacΔ(lacZ)M15 Δ(lacZYAargF)U169 zdg-232::Tn10 uidA::pir+* | Thermo Scientific, USA |

**F.** Constructed strains for *in vivo* adaptation assay.

| **Name** | **Bacterial strain** | **Plasmid name** | **Notes** | **Reference** |
|---|---|---|---|---|
| #965 | *E. coli* MG1655 | 964_p15A-IntI1$^{wt}$-smR2-catB3-blaVIM4-aac(6')-Ib-cr | IntI1$^{wt}$ | This study |
| #1010 | *E. coli* MG1655 | 977_p15A-IntI1$^{Y312F}$-smR2-catB3-blaVIM4-aac(6')-Ib-cr | IntI1$^{Y312F}$ | This study |
| #1016 | *E. coli* MG1655 | 1015_p15A-IntI1$^{\Delta C}$-smR2-catB3-blaVIM4-aac(6')-Ib-cr | IntI1$^{\Delta C}$ | This study |
| #1008 | *E. coli* MG1655 ΔlacZ::mCherry-pS-lox71/66 | 964_p15A-IntI1$^{wt}$-smR2-catB3-blaVIM4-aac(6')-Ib-cr | IntI1$^{wt}$ | This study |
| #1009 | *E. coli* MG1655 ΔlacZ::mCherry-pS-lox71/66 | 977_p15A-IntI1$^{Y312F}$-smR2-catB3-blaVIM4-aac(6')-Ib-cr | IntI1$^{Y312F}$ | This study |

**Table S4. Protein and peptide sequences used in this study. (A)** Full protein sequences of purified IntI1$^{Y312F}$ and IntI4$^{Y302F}$. **(B)** Peptide sequences. **(C)** IntI1 integrase sequences from the integron plasmid p15A, with indicated mutations. **(D)** mCherry-pS sequence.

**A.** Protein sequences IntI1$^{Y312F}$ and IntI4$^{Y302F}$ used for optical tweezers experiments. MBP tag is indicated in blue, integrase is indicated in yellow.

>MBP-IntI1$^{Y312F}$

MKIEEGKLVIWINGDKGYNGLAEVGKKFEKDTGIKVTVEHPDKLEEKFPQVAATGDGPDIIFWAHDRFGGYAQSGLLAEITPDK
AFQDKLYPFTWDAVRYNGKLIAYPIAVEALSLIYNKDLLPNPPKTWEEIPALDKELKAKGKSALMFNLQEPYFTWPLIAADGGY
AFKYENGKYDIKDVGVDNAGAKAGLTFLVDLIKNKHMNADTDYSIAEAAFNKGETAMTINGPWAWSNIDTSKVNYGVTVLPTFK
GQPSKPFVGVLSAGINAASPNKELAKEFLENYLLTDEGLEAVNKDKPLGAVALKSYEEELVKDPRIAATMENAQKGEIMPNIPQ
MSAFWYAVRTAVINAASGRQTVDEALKDAQTNSSSNNNNNNNNNNLGIEGRISHMSMGGRDIGRSENLYFQGSMKTATAPLPPL
RSVKVLDQLRERIRYLHYSLRTEQAYVHWVRAFIRFHGVRHPATLGSSEVEAFLSWLANERKVSVSTHRQALAALLFFYGKVLC
TDLPWLQEIGRPRPSRRLPVVLTPDEVVRILGFLEGEHRLFAQLLYGTGMRISEGLQLRVKDLDFDHGTIIVREGKGSKDRALM
LPESLAPSLREQLSRARAWWLKDQAEGRSGVALPDALERKYPRAGHSWPWFWVFAQHTHSTDPRSGVVRRHHMYDQTFQRAFKR
AVEQAGITKPATPHTLRHSFATALLRSGYDIRTVQDLLGHSDVSTTMIFTHVLKVGGAGVRSPLDALPPLTSER*

>MBP-IntI4$^{Y302F}$

MKIEEGKLVIWINGDKGYNGLAEVGKKFEKDTGIKVTVEHPDKLEEKFPQVAATGDGPDIIFWAHDRFGGYAQSGLLAEITPDK
AFQDKLYPFTWDAVRYNGKLIAYPIAVEALSLIYNKDLLPNPPKTWEEIPALDKELKAKGKSALMFNLQEPYFTWPLIAADGGY
AFKYENGKYDIKDVGVDNAGAKAGLTFLVDLIKNKHMNADTDYSIAEAAFNKGETAMTINGPWAWSNIDTSKVNYGVTVLPTFK
GQPSKPFVGVLSAGINAASPNKELAKEFLENYLLTDEGLEAVNKDKPLGAVALKSYEEELVKDPRIAATMENAQKGEIMPNIPQ
MSAFWYAVRTAVINAASGRQTVDEALKDAQTNSSSNNNNNNNNNNLGIEGRISHMSMGGRDIGRSENLYFQGSMKSQFLLSVRE
FMQTRYYAKKTIEAYLHWITRYIHFHNKKHPSLMGDKEVEEFLTYLAVQGKVATKTQSLALNSLSFLYKEILKTPLSLEIRFQR
SQLERKLPVVLTRDEIRRLLEIVDPKHQLPIKLLYGSGLRLMECMRLRVQDIDFDYGAIRIWQGKGGKNRTVTLAKELYPHLKE
QIALAKRYYDRDLHQKNYGGVWLPTALKEKYPNAPYEFRWHYLFPSFQLSLDPESDVMRRHHMNETVLQKAVRRSAQEAGIEKT
VTCHTLRHSFATHLLEVGADIRTVQEQLGHTDVKTTQIFTHVLDRGASGVLSPLSRL*

**B.** Peptide sequences.

| Name | Sequence | Mimicking IntI | Notes |
|---|---|---|---|
| pL | RSPLDALPPLTSER | IntI1 | Full tail |
| pS | RSPLDAL | IntI1 | Only α-helix |
| pS‘ | R**A**PLDAL | IntI1 | Only α-helix, mutated |
| pS“ | RSP**A**DAL | IntI1 | Only α-helix, mutated |
| pS4 | LSPLSRL | IntI4 | Full tail |

**C.** IntI1 integrase sequences from the integron plasmid p15A, with indicated mutations.

> IntI1$^{wt}$

MKTATAPLPPLRSVKVLDQLRERIRYLHYSLRTEQAYVHWVRAFIRFHGVRHPATLGSSEVEAFLSWLANERKVSVSTHRQALAA
LLFFYGKVLCTDLPWLQEIGRPRPSRRLPVVLTPDEVVRILGFLEGEHRLFAQLLYGTGMRISEGLQLRVKDLDFDHGTIIVREG
KGSKDRALMLPESLAPSLREQLSRARAWWLKDQAEGRSGVALPDALERKYPRAGHSWPWFWVFAQHTHSTDPRSGVVRRHHMYDQ
TFQRAFKRAVEQAGITKPATPHTLRHSFATALLRSGYDIRTVQDLLGHSDVSTTMIYTHVLKVGGAGVRSPLDALPPLTSER*

| |
|---|
| > IntI1$^{Y312F}$<br><br>MKTATAPLPPLRSVKVLDQLRERIRYLHYSLRTEQAYVHWVRAFIRFHGVRHPATLGSSEVEAFLSWLANERKVSVSTHRQALAA<br>LLFFYGKVLCTDLPWLQEIGRPRPSRRLPVVLTPDEVVRILGFLEGEHRLFAQLLYGTGMRISEGLQLRVKDLDFDHGTIIVREG<br>KGSKDRALMLPESLAPSLREQLSRARAWWLKDQAEGRSGVALPDALERKYPRAGHSWPWFWVFAQHTHSTDPRSGVVRRHHMYDQ<br>TFQRAFKRAVEQAGITKPATPHTLRHSFATALLRSGYDIRTVQDLLGHSDVSTTMI**F**THVLKVGGAGVRSPLDALPPLTSER* |
| > IntI1$^{\Delta C}$<br><br>MKTATAPLPPLRSVKVLDQLRERIRYLHYSLRTEQAYVHWVRAFIRFHGVRHPATLGSSEVEAFLSWLANERKVSVSTHRQALAA<br>LLFFYGKVLCTDLPWLQEIGRPRPSRRLPVVLTPDEVVRILGFLEGEHRLFAQLLYGTGMRISEGLQLRVKDLDFDHGTIIVREG<br>KGSKDRALMLPESLAPSLREQLSRARAWWLKDQAEGRSGVALPDALERKYPRAGHSWPWFWVFAQHTHSTDPRSGVVRRHHMYDQ<br>TFQRAFKRAVEQAGITKPATPHTLRHSFATALLRSGYDIRTVQDLLGHSDVSTTMIYTHVLKVGG** |

**D**. mCherry-pS sequence. 8xHis-tag is indicated in orange, mCherry is indicated in red and the pS peptide is violet.

| |
|---|
| > 8xHis-mCherry-pS<br><br>HHHHHHHHGSSGMVSKGEEDNMAIIKEFMRFKVHMEGSVNGHEFEIEGEGEGRPYEGTQTAKLKVTKGGP<br>LPFAWDILSPQFMYGSKAYVKHPADIPDYLKLSFPEGFNWERVMNFEDGGVVTVTQDSSLQDGEFIYKVK<br>LRGTNFPSDGPVMQCRTMGWEASTERMYPEDGALKGEIKQRLKLKDGGHYDAEVKTTYKAKKPVQLPGAY<br>NVDIKLDILSHNEDYTIVEQYERAEGRHSTGGMDELYKSENLYFQGVRSPLDAL* |

**Table S5. Python 3.10.5 packages used in this study.**

| | |
|---|---|
| lumicks.pylake 1.2.0 | pathlib |
| numpy 1.26.3 | pandas 2.1.4 |
| glob | scipy 1.11.4 |
| math | seaborn 0.12.2 |
| statistics | pingouin 0.5.3 |
| os | statsmodels 0.14.0 |
| matplotlib 3.8.0 | |
